# Functional advantages of Lévy walks emerging near a critical point

**DOI:** 10.1101/2020.01.27.920801

**Authors:** Masato S. Abe

## Abstract

A special class of random walks, so-called Lévy walks, has been observed in a variety of organisms ranging from cells, insects, fishes, and birds to mammals, including humans. Although their prevalence is considered to be a consequence of natural selection for higher search efficiency, some findings suggest that Lévy walks might also be epiphenomena that arise from interactions with the environment. Therefore, why they are common in biological movements remains an open question. Based on some evidence that Lévy walks are spontaneously generated in the brain and the fact that power-law distributions in Lévy walks can emerge at a critical point, we hypothesized that the advantages of Lévy walks might be enhanced by criticality. However, the functional advantages of Lévy walks are poorly understood. Here, we modeled nonlinear systems for the generation of locomotion and showed that Lévy walks emerging near a critical point had optimal dynamic ranges for coding information. This discovery suggested that Lévy walks could change movement trajectories based on the magnitude of environmental stimuli. We then showed that the high flexibility of Lévy walks enabled switching exploitation/exploration based on the nature of external cues. Finally, we analyzed the movement trajectories of freely moving *Drosophila* larvae and showed empirically that the Lévy walks may emerge near a critical point and have the large dynamic range and high flexibility. Our results suggest that the commonly observed Lévy walks emerge near a critical point and could be explained on the basis of these functional advantages.

## Introduction

Lévy walks are a special class of random walks with step lengths ℓ that follow a power-law distribution *P* (ℓ) ~ ℓ^−*μ*^ where *μ* ∈ (1, 3] is a power-law exponent. Lévy walks are observed in a variety of biological movements and agents, ranging from cells and insects to mammals, humans, and even memory retrievals in human cognition [1, 2, 3, 4, 5, 6, 7, 8]. Lévy walks are composed of many short steps and rare, long straight movements. This characteristic leads to more efficient searching or foraging strategies than normal random walks, so-called Brownian walks, when the targets (e.g., food, mates, or habitats) are unpredictably and sparsely distributed in the environment [9, 10, 11, 12]. According to the Lévy walk foraging hypothesis, evolution through natural selection can therefore explain why Lévy walks are widespread in biological movements.

However, there has been a recent debate about the origin of Lévy walks. Some findings suggest that Lévy walks might also be epiphenomena arising from interactions with a complex environment [13, 14]. For example, the Lévy walks observed in the movements of plant seeds are thought to result from interactions between wind turbulence and seed shape. Thus, why Lévy walks are a common mode of biological movement and how they are generated remain controversial. One of the causes of this controversy is uncertainty about the mechanism (e.g., stochastic decisions and environment) that generates Lévy walks and their advantages (e.g., foraging efficiency). Understanding why Lévy walks are prevalent in biological movements therefore requires determining how Lévy walks emerge based on biological mechanisms and what functional advantages they provide.

One of the candidate mechanisms responsible for Lévy walks is the spontaneous dynamics generated by internal states such as nervous systems. Previous studies on spontaneous behavior have revealed that the temporal patterns of the intervals between consecutive turns or rest/movement without a stimulus often follow a power-law distribution [15, 16, 17], which is a hallmark of Lévy walks. In addition, the frequent correspondence of movement trajectories of some animal species in a featureless environment to Lévy walks has suggested that Lévy walks are generated intrinsically [18, 19]. Moreover, the revelation by a recent study that *Drosophila* larvae exhibit Lévy walks even when their sensory neurons are blocked is strong evidence that Lévy walks are intrinsically generated, especially in the central pattern generators (CPGs) of the brain [20].

Such temporal organizations of spontaneous behavior or movement patterns in Lévy walks in some species could be based on nonlinear dynamics or chaotic dynamics [21, 15, 22, 23]. Theoretical studies have shown that low-dimensional dynamical systems can produce Lévy walks [24, 25]. Furthermore, power-law distributions are often observed in a system close to a critical point between order and disorder [26]. Importantly, being near a critical point can lead to functional advantages such as optimal dynamic ranges, high sensitivity, and computational ability, and so on [27, 26]. In particular, optimal dynamic ranges provide the ability to code information spanning several orders of magnitude, which is essential for biological systems to respond to physical stimuli [27]. These findings suggest that Lévy walks emerging near a critical point may benefit from criticality. However, what advantages they may confer to biological agents remain poorly understood.

In this study, we explored the functional advantages of Lévy walks emerging near a critical point by using a model composed of simple, deterministic, nonlinear systems for generating movements. We found that Lévy walks near a critical point can have a large dynamic range for environmental stimuli. This large dynamic range implies that a system producing Lévy walks near a critical point can functionally process information and can change the movement patterns in response to several orders of stimuli. Moreover, the ability of Lévy walks to switch between exploration and exploitation based on the kind of sensory information suggests that Lévy walks provide an advantage that enables responses adapted to environmental conditions. Finally, we analyzed the movement trajectories of *Drosophila* larvae and confirmed that their Lévy walks could emerge near a critical point and had a large dynamic range and high flexibility that were consistent with our theoretical models. These findings provide novel insights into understanding how and why Lévy walks emerge in biological systems.

## Results

### Movements obtained from the model

We modeled the movements of autonomous agents (e.g., individual animals) based on their internal states. The CPGs in brains have been shown to produce locomotion patterns even without external stimuli [28, 29, 30]. Because CPGs are composed of coupled oscillators [28, 29], it is reasonable to construct a minimal model composed of a few (two in the main text) nonlinear oscillators, *x_t_* and *y_t_*, of the agents (see Materials and Methods), as pointed out by [31]. In *Drosophila* larvae, there are two neural circuits, the thoracic and abdominal networks, that produce search behavior [30]. Their activities can thus be considered to correspond to *x_t_* and *y_t_*. Because the signature of chaos has been found in movements and neuronal firings [22, 32, 33], we required the nonlinear function of the model to be chaotic dynamics (here, a tent map for tractability). Moreover, because connections between elements are ubiquitous in biological systems, the model included a parameter *ε* ∈ [0, 0.5] that described the coupling (connection) strength between elements in the system. For example, *ε* corresponded to the strength of interaction between neural units. We then considered the motor outputs from the dynamics, *x_t_* and *y_t_* (Fig. 1A). In the CPGs of some taxa, some neuronal units are connected to right and left motor outputs [28]. In other cases, the two variables could be the activities of two neural circuits in CPGs. For example, when the thoracic and abdominal networks in *Drosophila* larvae have symmetrical outputs, they exhibit straight-line crawls. In contrast, when the outputs are asymmetric, they exhibit turnings [30]. Based on these observations, we assume that movement is generated from the turning angle Δ*θ_t_* = *c*(*x_t_* – *y_t_*) of the movements at each time *t* (Fig. 1A), where *c* is a constant that connects internal dynamics and outputs (see Materials and Methods). When the two outputs are symmetric, the step corresponds to a straight motion without turns. For simplicity, we assume that the speed of movement is constant. Note that we do not assume any distributions of step lengths in the movements and turning angles.

**Figure 1:**
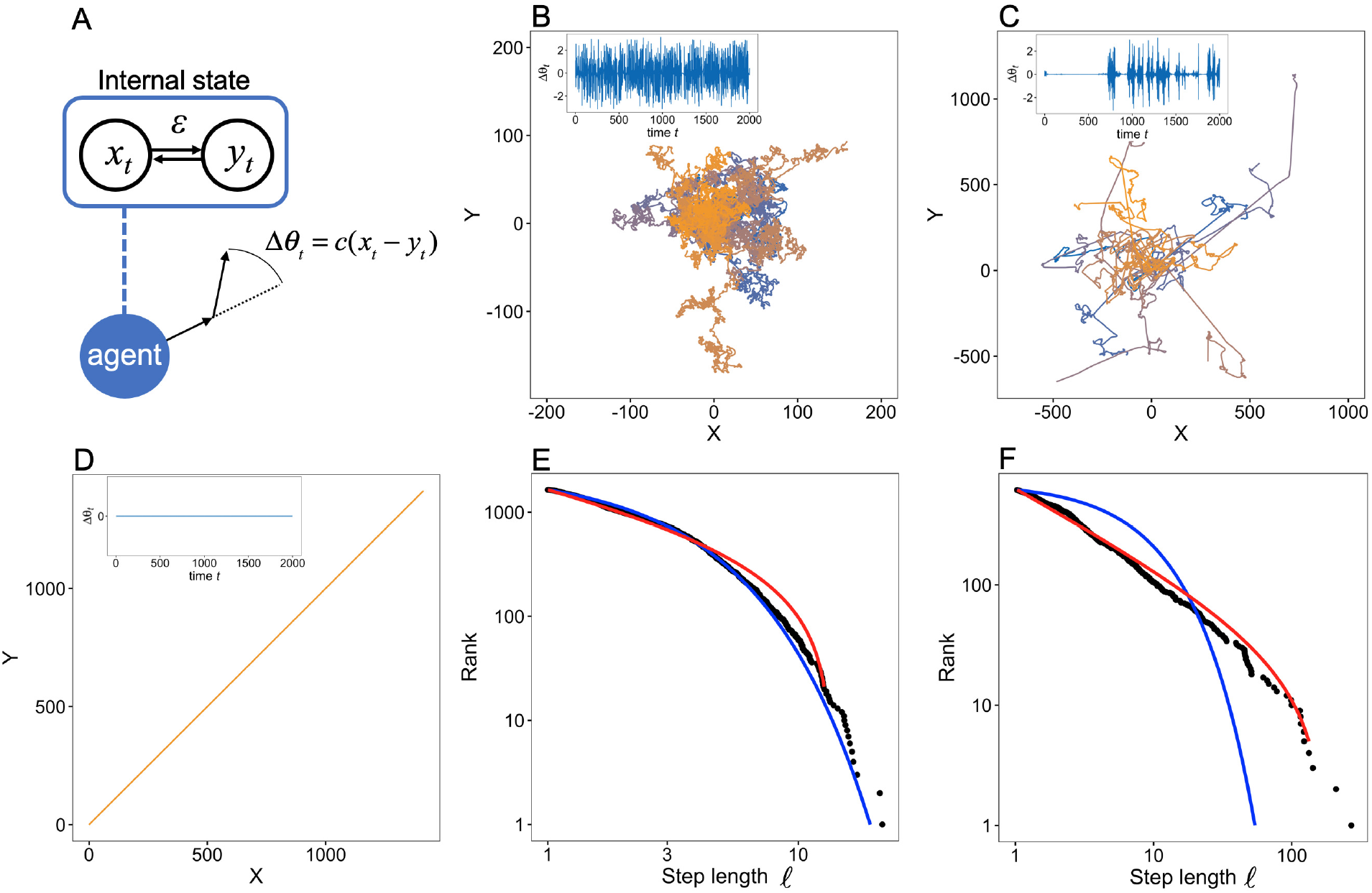
Our model scheme and examples of trajectories and step length distributions obtained from the model. (A) The internal dynamics *x_t_* and *y_t_* produce the agent movements. The model has a parameter that determines the coupling strength between elements in the system. The movement is simply produced by turning angles Δ*θ_t_*. (B-D) The trajectory of the agent in a two-dimensional space from *t* = 0 to 2,000 is represented by solid lines: (B) *ε* = 0.1; (C) *ε* = 0.22; (D) *ε* = 0.3. The different colors correspond to different initial conditions. Note that the trajectories in (D) are overlaid because they are exactly the same. The parameter of the tent map was set to *r* = 0.7, and the initial position ***X*** (0) was set to (0, 0). Examples of the time series of the turning angle Δ*θ_t_* are shown in the insets. When the value of Δ*θ_t_* is close to zero for a long time, the agent exhibits straight movement. In contrast, when Δ*θ_t_* fluctuates dynamically, the agent makes many turns. (E) and (F) Log-log plots of the rank distribution of step lengths ℓ in the trajectories obtained from the model for *ε* = 0.1 and *ε* = 0.22, respectively. Black dots represent the distribution of step length from the model movements as shown in (B) and (C). The other parameters were set to *r* = 0.7 and *t*_max_ = 10,000. The red and blue lines represent the fitted distribution of the truncated power-law and the exponential distribution, respectively.

First, we show some examples of movement trajectories of agents obtained from the model and the dependence of those trajectories on the control parameter *ε* (Fig. 1B-D). When *ε* = 0.1—namely, the components of the systems interacted weakly with each other—the movement was similar to Brownian walks (Fig. 1B). This similarity reflected the fact that the two chaotic oscillators had less influence on each other, and the turning angles could randomly fluctuate as if they were independent random numbers. In contrast, when *ε* = 0.3, the movement was completely ballistic—namely, a straight line (Fig. 1D)—because the two oscillators were completely synchronized (*x_t_* = *y_t_*) due to the strong interaction strength. Hence *x_t_* and *y_t_* fluctuated in the same manner. At an intermediate level of interaction strength, here *ε* = 0.22, it is apparent that the movement was composed of rare, long, straight movements among many short steps (Fig. 1C). This pattern is characteristic of Lévy walks. The difference of the movement trajectories between *ε* = 0.1 and 0.22 was also reflected in the distribution of step lengths (Figs. 1E and F). A rigorous fitting method and the evaluation of the diffusive properties supported the conclusion that the movement trajectories for *ε* = 0.1 and 0.22 were Brownian walks and Lévy walks, respectively (see Fig. S1 and Materials and Methods) [34, 35, 36].

An illustration of the internal dynamics and the attractors corresponding to the movements makes it possible to intuitively grasp the mechanisms that produce such movements (insets of Figs. 1B–D and S2). Note that Δ*θ_t_* in the insets of Fig. 1B-D is proportional to *x_t_* – *y_t_*, and thus Δ*θ_t_* = 0 means that *x_t_* and *y_t_* are synchronized with each other. The internal dynamics that produced the Lévy walk-like movements exhibited intermittent dynamics (inset of Fig. 1C). This pattern reflects the fact that intermittent searches with fractal reorientation clocks correspond to Lévy walks [37]. In contrast, the dynamics were always noisy for small *ε* (inset of Fig. 1B) and were entirely absent for large *ε* (inset of Fig. 1D). These results indicate that the various movement patterns arose from the synchronous and asynchronous dynamics of the internal states.

### Phase diagram

To elucidate the conditions that are needed for the emergence of Lévy walks in our model, we then drew a phase diagram (Fig. 2) by changing the control parameter *ε* and evaluating the internal dynamics and movement patterns. The trajectories for the same parameter set were generated by different initial conditions *x*_0_ and *y*_0_ (100 replicates). To characterize the internal dynamics, the order parameter of the system was equated to the standard deviation of the distributions of Δ*θ* averaged over 100 replicates. The demonstration in Fig. 2 that the order parameter changed drastically from positive standard deviations to zero standard deviations for *ε* ≈ 0.23 suggested that the system exhibited a phase transition. This was also a critical point at which the synchronized state *x_t_* = *y_t_* changed from an unstable state to a stable one with increasing *ε*. This transition can be characterized by the linear stability of *x_t_* – *y_t_* (see Supplementary Information).

**Figure 2:**
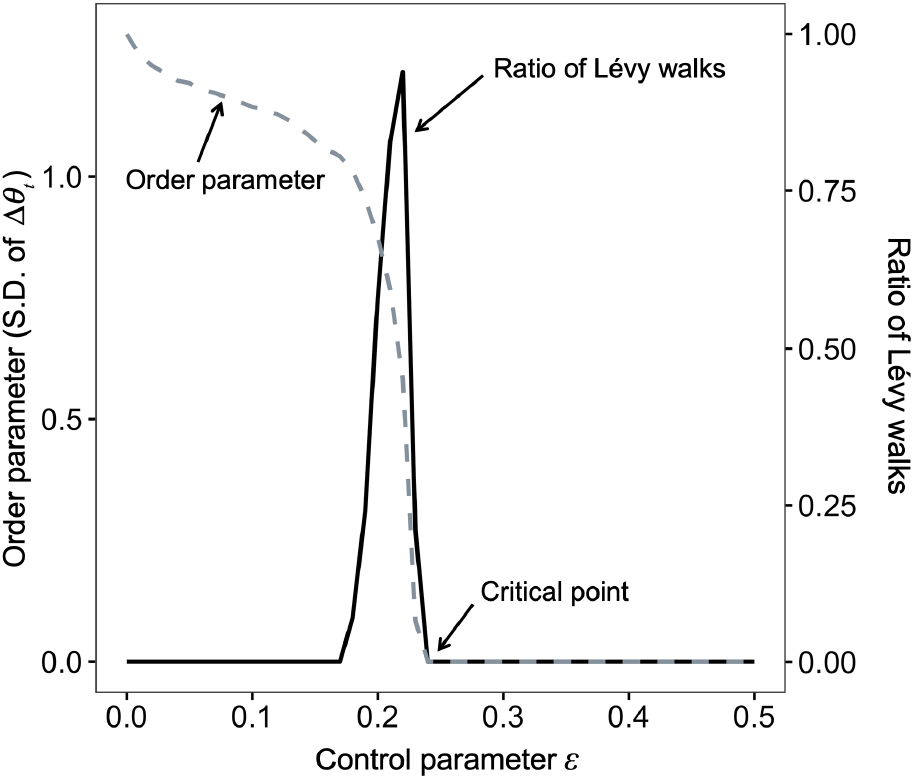
Phase diagram with a changing control parameter *ε*. The gray dashed line represents the order parameter, which is the standard deviation of the distribution of Δ*θ_t_*. Near *ε* = 0.22 the order parameter changes drastically. The solid black line represents the ratio of the classification of Lévy walks obtained by fitting distributions to the simulated 100 trajectories for the same parameter set. The other parameters were set to *r* = 0.7 and *t*_max_ = 10,000.

We also conducted a classification of the movements obtained with different *ε* and initial conditions using statistical techniques as well as the above section. The results showed that the region of emergence of Lévy walks was poised near the critical point between the synchronous and asynchronous phases, which correspond to straight movements and Brownian walks, respectively (Fig. 2). At *ε* = 0.22, the ratio of walks was maximized, and the estimated power-law exponent was 1.55 ± 0.06 (mean ± S.D.).

The transition occurred via intermittent patterns, as shown in the inset of Fig. 1C, which is called a blowout bifurcation [38, 39, 40]. Near the critical point of the bifurcation, log(|*x_t_* – *y_t_*|) exhibited unbiased random walks. According to random walk theory, the return time from a value to the same value follows a power-law distribution. Thus, |*x_t_* – *y_t_*| could exhibit intermittent behavior [40].

Because the search efficiency depends on the distribution of step lengths and the diffusiveness [9, 10], the Lévy walkers developed by our model also received the benefits of a search efficiency higher than that achieved with Brownian walks. In addition, it is worth noting that the trajectories could change as functions of the initial internal states, *x*_0_ and *y*_0_, while keeping the characteristics of the movement patterns mostly in the parameter range (Figs. 1B–D and Fig. S3). This possibility can explain the variability that is often observed in animal decision making [15, 41].

Moreover, we confirmed that the emergence of Lévy walks near a criticality is apparent in other models, including a model based on different *r* or logistic maps (i.e., different nonlinear function form *f*) (Figs. S5 and S6) [42], global coupled maps (i.e., many components) (Fig. S7) [43], and noise-additive models (Fig. S8).

### Optimal dynamic range

Once we had modeled the Lévy walks emerging near a critical point based on biological assumptions, we explored the functional advantages, except for searching efficiency under uninformed conditions. The demonstration that some complex systems sitting near a critical point can have large dynamic range means that they can code information spanning several orders of magnitude [27, 26]. This characteristic is an advantage for information processing, and thus we expect that a model exhibiting Lévy walks will also have a large dynamic range.

To investigate dynamic ranges, we added perturbations (i.e., stimuli) of different amplitudes *S* to the internal dynamics and then observed the movement trajectories (Fig. 3A; see the details in Materials and Methods). First, we evaluated the response *F* as the distance between the positions of the agent with and without the perturbation. Fig. 3B shows the dependence of *F* on *S*. In the region below the critical point (i.e., *ε* < 0.22), *F* was insensitive to perturbations because the stimuli were masked by the highly spontaneous, variable activities. In contrast, although in the region above the critical point (*ε* > 0.23) *F* could change in response to stimuli, it could respond only to large stimuli. Near the critical point, *F* could change in response to stimuli several orders of magnitude in size. To evaluate these characteristics, we calculated the dynamic range Δ*_S_* (see the inset of Fig. 3C and Materials and Methods). Fig. 3C shows that the dynamic range was largest near the critical point for various *τ*. This result suggested that Lévy walks had an optimal dynamic range and could respond to the stimuli of several orders of magnitude in size. These characteristic imply that systems producing Lévy walks near a critical point have information about stimuli and can functionally process that information. In addition, Lévy walks have large *F* for most *S* (Fig. 3B). This indicates that an agent can have high sensitivity to the stimuli and drastically change its movements. In addition to having an optimal dynamic range, an agent at a critical point has the ability to change its movement in response to stimuli. For example, when an agent obtains novel information about the presence of new resources such as food or habitat, it should change its originally planned movement trajectory. In another example, when an agent receives a stimulus about the presence of predators, it should avoid them by changing its original movement trajectory according to the intensity of the signal. Our results suggested that Lévy walks emerging near a critical point could deal with these situations.

**Figure 3:**
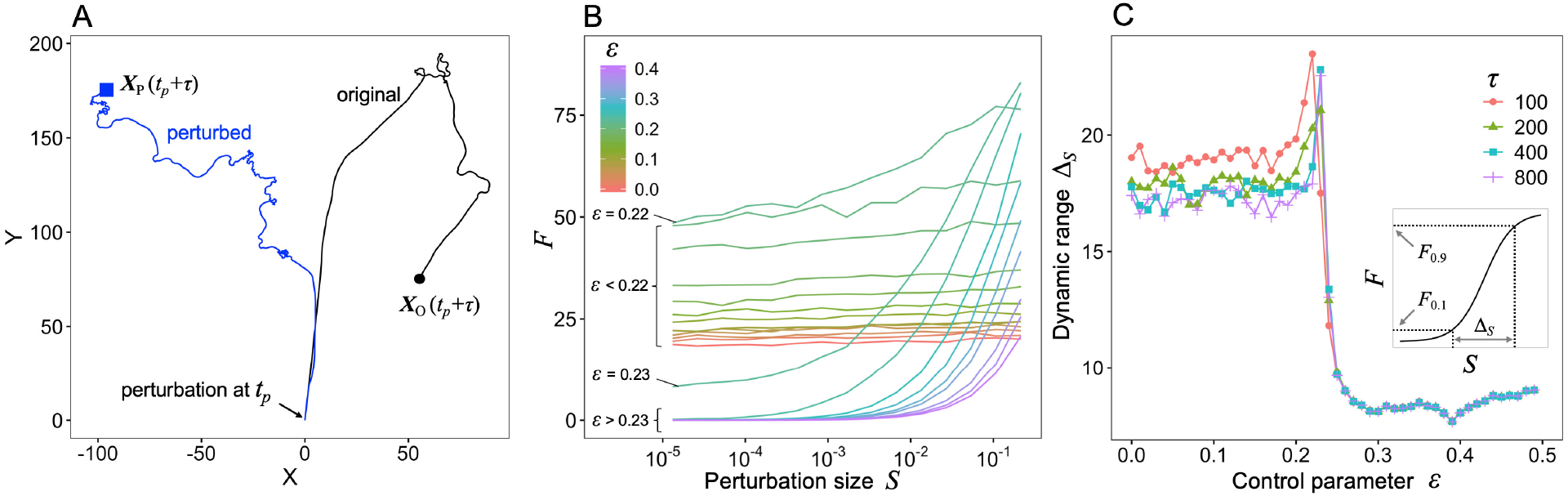
Dynamic range (A) An example of trajectories for *ε* = 0.22 with (blue line) and without (black line) perturbations. We added a perturbation *S* = 10^−4^ to the internal dynamics of *x* and *y* only at *t_p_* and ***X*** (*t_p_*) = (0, 0). The black circle and blue square represent the positions at *t* = *t_p_* + *τ* (here *τ* = 400) of the original and perturbed trajectories, respectively. A trajectory change *F* is defined as the distance between ***X***_O_ (*t_p_* + *τ*) and ***X***_P_ (*t_p_* + *τ*). (B) Relationship between perturbation size *S* and *F* for *τ* = 100. (C) Control parameter *ε* and dynamic range Δ*_S_*. The inset represents the definition of dynamic range Δ*_S_*. These results represent values averaged over 10^3^ simulation runs with different initial conditions.

### Adaptive switching between exploitation and exploration

Whereas uninformed animals should exhibit random searches such as Lévy walks under many conditions [1], animals also need to switch between exploitation and exploration in response to sensory information about the environment [44]. For instance, when they obtain information about the presence of targets close to them, they may undertake an area-restricted search for the nearby place (i.e., exploitation). In contrast, when they obtain information about the absence of targets, they may give up searching for the nearby place and look for a new place (i.e., exploration). Such switching based on sensory information allows searchers to obtain more targets [45].

To reveal the relationship between adaptive switching and criticality, we analyzed the dependence of movement patterns on a stimulus parameter *S*′ (see Material and Methods). The distance *G* between the position where the agent received a stimulus and the position after time *τ* was calculated to quantify the degree of change of movements, namely switching between exploitation and exploration (see Fig. 4A). A small *G* means that the agents, after receiving the stimulus, stayed near the place where they received the stimulus, and vice versa. Therefore, when many targets are near the agent, a small *G* is a better strategy (i.e., exploitation), whereas a large *G* is a better strategy (i.e., exploration) when there are no targets near the agent. Fig. 4B shows the dependence of *G* on the stimulus parameter *S*′. Whereas the *G* for Brownian walks (small *ε*) and straight movements (*ε* > 0.22) was less sensitive to stimuli, the *G* for Lévy walks (*ε* ~ 0.22) could respond to a wide variety of stimuli. As we did in the previous section, we defined the dynamic range of *G* (see details in the inset of Fig. 4C and Materials and Methods). The large dynamic range of agents near the critical point (Fig. 4C) suggested that Lévy walks emerging near a critical point could change the movement patterns according to information about the type of stimuli (e.g., amount of food). We also investigated the response range Δ*_G_* = *G*_max_ – *G*_min_ to determine how much the response of an agent could vary as a function of the stimulus parameter. The indication in Fig. 4D that agents near the critical point have a large response range suggested that Lévy walks could flexibly switch between exploration and exploitation based on the types of stimuli they receive. This flexibility reflects the fact that the internal dynamics fluctuates intensely when *ε* is below the critical point (inset of Fig. 1B), and these fluctuations cause numerous turns, irrespective of *S*′. In contrast, the stable internal dynamics for *ε* above the critical point produces straight movements (i.e., few turns), even in the presence of large *S*′. Movements near the critical point have the potential ability to switch between exploitation and exploration thanks to the small positive Lyapunov exponents.

**Figure 4:**
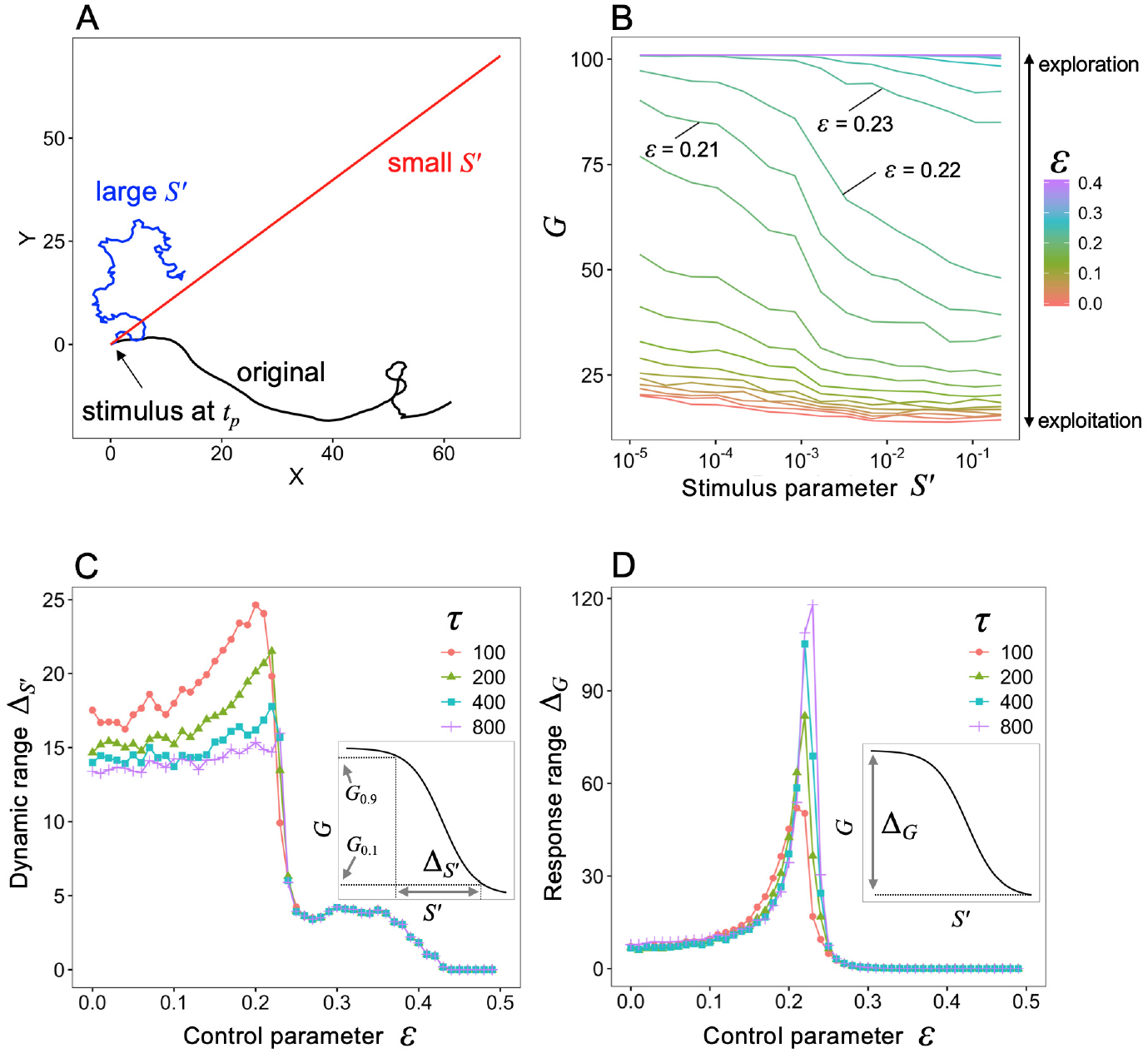
Switching between exploitation and exploration (A) An example of trajectories for *ε* = 0.22 with (red, *S*′ = 10^−4^; blue, *S*′ = 0.3) and without (black) stimuli. The agents receieved the stimuli only at *t_p_* and *X*(*t_p_*) = (0, 0). We observed the position after *τ* (here *τ* = 100) and calculated the distance *G* between *X*(*t_p_*) = (0, 0) and *X*(*t_p_* + *τ*). (B) Relationship between stimuli parameters *S*′ and *G* for *τ* = 100. (C) Dynamic range Δ*_S′_* and control parameter *ε*. The inset represents the definition of Δ*_S′_*. (D) Response range Δ*_G_* and control parameter *ε*. The inset represents the definition of Δ*_G_*. These results represent values averaged over 10^3^ simulation runs with different initial conditions.

### Verification with empirical data

To verify our model, we used empirical datasets of movements of *Drosophila* larvae exhibiting Lévy walks from [20], as shown in Fig. 5A. The exponents of the truncated power-law distributions averaged 1.59 ± 0.45 (mean ± S.D., *n* = 304), which is similar to the exponent of 1.55 ± 0.06 in the present model with *ε* = 0.22 (see Phase diagram section).

**Figure 5:**
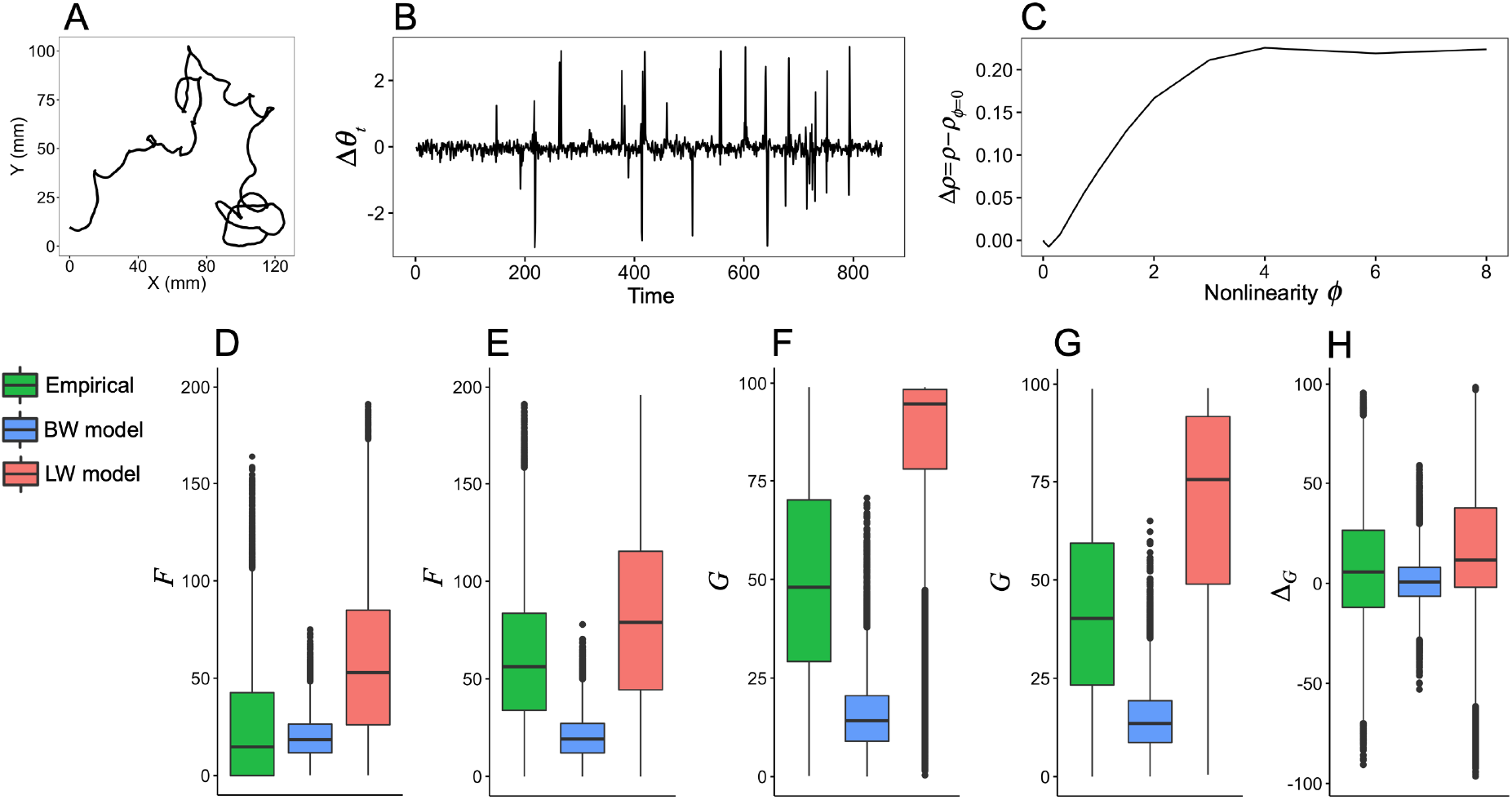
Empirical anlaysis. (A) and (B) Example of a movement trajectory of *Drosophila* larvae and the corresponding time series of turning angles, respectively. (C) Example of the result of S-map to check nonlinearity of the time series. (D-H) The results of perturbation analysis. (D) and (E) The corresponding cases of small and large *S* for *τ* = 100, respectively. (F) and (G) The corresponding cases of small and large *S*′ for *τ* = 100, respectively. (H) The difference of *G* between (F) and (G). Note that BW and LW model mean the proposed model with *ε* = 0.0 and 0.22, respectively. In (D-H), all pairwise differences among the empirical, BW model, and LW model were statistically significant (*p* < 10^−15^). The black dots in (D-H) represent outliers, defined as data points outside the whiskers, which correspond to 1.5 times the interquartile range.

We first investigated whether the time series of turning angles Δ*θ_t_* (Fig. 5B) was generated from nonlinear systems or stochastic processes. To make this determination, we used a nonlinear prediction method with a parameter *ϕ* controlling the nonlinearity of the prediction model (see the details in Materials and Methods) [46, 47]. If the time series was generated from nonlinear systems, the prediction accuracy *ρ* would be higher for the nonlinear prediction (*ϕ* > 0) than the linear one (*ϕ* = 0) (Fig. 5C). We found that the nonlinear predictions were better than the linear ones in 86.3% of the time series (251 of 291). Moreover, the differences between the nonlinear and linear predictions were found to be statistically significant in 48.1% of the time series (140 of 291). These results suggested that most of the time series were generated through nonlinear systems rather than via stochastic processes.

Once we had confirmed the nonlinearity of the turning angles, we calculated the largest Lyapunov exponents of these time series and found that they were 0.13 ± 0.15 (mean ± S.D.) (Fig. S9). In addition, the largest Lyapunov exponents of time series obtained from the proposed model with *ε* = 0.0 and 0.22 were 0.32 ± 0.14 and 0.07 ± 0.04, respectively. This result suggested that some of the empirical time series could be generated from a system near a critical point.

Finally, we analyzed how perturbations could change the movement trajectories of empirical data. To carry out this analysis, we conducted a data-based simulation based on a twin surrogate algorithm (see details in Materials and Methods and Supplementary Information) [48]. For comparison, we also conducted the same analyses on time series obtained from the present model (*ε* = 0.0 and 0.22). Fig. 5D and 5E show the response *F* for the cases of small and large *S*, respectively. In the empirical data and the Lévy walk model, the *F* for large perturbations was substantially larger than the *F* for small perturbations. This difference suggests that Lévy walks by *Drosophila* larvae have large dynamic ranges as well as the Lévy walk model. Figs. 5F and 5G show the response *G* for the cases of small and large *S*′, respectively. Whereas the Brownian walks model was insensitive to perturbation size, the *G* of the empirical data could change in response to the stimulus parameter (Figs. 5F and G). This flexibility is reflected in Fig. 5H, which shows that the empirical Δ*_G_* (difference of *G* between the cases of small and large *S*′) was larger than the Δ*_G_* for the Brownian walk model. This suggests that the Lévy walks in *Drosophila* larvae can also change movement patterns. These results are qualitatively consistent with the predictions from our theoretical analysis. The difference of *F* and *G* between the empirical movements and the Lévy walk model observed in Figs. 5D–H can be attributed to the difference of the diffusion property. The mean exponent *α* of the MSD ~ *t^α^* for each group condition in the empirical data ranged from 0.94 to 1.41 [20], whereas the exponent of the Lévy walk model was 1.73 up to *t* ≈ 500 (Fig. S1). It is thus plausible that a Lévy walk model with a higher diffusivity will have higher *F* and *G*.

## Discussion

In the present study, we modelled Lévy walks based on dynamical systems such as CPGs to address the questions of why, in terms of functional advantages, animals often exhibit Lévy walks. We showed that Lévy walks emerging near a critical point can have large dynamic ranges and be highly flexible (Figs. 3 and 4). In addition, we confirmed these results by using empirical movement data. Based on these results, we propose that Lévy walks are commonplace because of the functional advantages they derive from being near a critical point.

The mechanisms that generate biological Lévy walks can be classified into two types: mechanisms based on interaction with environments and those based on internal states. Although the former mechanisms are consistent with cases of Lévy walks observed in biological movements [14], the empirical evidence for intrinsically generated Lévy walks has been accumulating [18, 19, 20]. Theoretical studies on search efficiency have implicitly assumed that the movement patterns are caused by intrinsic mechanisms. However, those studies have focused only on the probability distribution of the step lengths and turning angles, and they have thus ignored the intrinsic generative mechanisms [9, 10, 12]. When one tries to fully understand why Lévy walks are widely observed in biological systems, it is essential to explore how the brain or systems related to behavior can produce Lévy walks intrinsically [31, 23, 49, 50]. Here, we assumed the simplest deterministic nonlinear model for intrinsically generated movements. Nevertheless, the produced movement patterns were complex enough to model the movements in real organisms and varied from Brownian walks to Lévy walks and ballistic movements, depending on the parameters (Figs. 1, 2, and S3). This variability reflects the synchronous and asynchronous dynamics of internal states that are ubiquitous in the brain [51]. Whereas animals with high cognitive ability may adopt Lévy walks based on rational decision-making [49], it is reasonable that animals with low cognitive ability, such as insects, adopt a simpler mechanism comprised of nonlinear chaotic oscillators, as shown in our empirical analysis (Fig. 5). The fact that only one parameter, the coupling strength, is required for controlling movement patterns in our model suggests that it is relatively easy to obtain and maintain Lévy walks through natural selection, in contrast to other mechanisms that involve many parameters.

It has been shown that power-law distributions, which are characteristic of Lévy walks, are often observed near a critical point in various systems. Examples include self-organized criticality of complex systems [52, 26] or excitable network models of the brain [27]. Furthermore, it has already been reported that deterministic chaos can lead to Lévy walks in the context of physics [24, 25]. As for search efficiency, the relationship between critical survival density and Lévy walks at the edge of extinction has been reported [53]. It should thus be emphasized that the idea of generative mechanisms based on criticality that we proposed here is not novel, although the relationship between biological Lévy walks based on internal states such as CPGs and criticality has not been elucidated.

In our model, the coupling strength in the system can drastically alter movement patterns (Figs. 1 and 2). Theoretical studies have shown that the optimal searching strategy can change as a function of the ecological context, such as the density of targets, risk of predation, and kinds of objects being sought [54, 55, 56, 57, 58]. Some empirical studies have reported that animals do not always exhibit Lévy walks [4, 5, 3]. It is therefore possible that the plasticity of spontaneous behavior caused by changing the coupling strength of the systems can lead to adaptive responses to environmental conditions. Moreover, differences in movement patterns can also impact not only the fitness of the agent but also higher systems, such as epidemic dynamics and ecosystems, through interactions with other agents. For instance, Lévy walks allow more resilient prey-predator systems than normal diffusion movements would under degraded conditions [59]. Understanding the mechanisms and controllability of Lévy walks can therefore lead to predicting and controlling the consequences in higher systems.

The main advantage of Lévy walks has previously been considered to be their high search efficiency compared to that of Brownian walks [1, 12, 13]. Whereas some studies have reported the robustness of the search efficiency of Lévy walks under various conditions [11, 12], a recent study has claimed that the optimal search strategy of Lévy walks with *μ* ≈ 2 in two or higher dimensions is not robust when non-destructive targets are randomly and sparsely distributed [58]. Based on this controversy and the dependence on ecological contexts of the diversity of optimal strategies, optimal foraging could not be the only strong evolutionary force that favors Lévy walks. Our results suggest that Lévy walks near the critical point outperform Brownian walks and ballistic movements within the model in terms of dynamic range and flexibility in changing behavior (Figs. 3–4). A large dynamic range makes it possible to distinguish the sizes of perturbations or stimuli in various kinds of information, such as amounts of food or presence of predators. Such information can lead to adaptive responses based on the environment. One such example is the adaptive switching shown in Fig. 4 that clearly led to obtaining more targets [45]. As another example, straight movements derived from a small *S*′ allow the agents to keep away from dangers such as predators. These results are consistent with the fact that optimal information processing can be found near a critical point [27, 26].

In the context of brain dynamics and complex systems, a critical point hypothesis has been proposed [60, 61, 26, 62, 63, 64]. The hypothesis claims that systems positioned at or near a critical point between ordered and disordered states have various functional advantages ranging from adaptive responses to a fluctuating environment to the production of various scale behaviors and computational abilities [26, 61, 65]. It is therefore reasonable that various biological systems, including brains, gene regulatory networks, and cell and animal groups that exhibit collective motions, carry the signature of being near a critical point [60, 26, 62, 63]. Likewise, in our model, agents with internal states near the criticality between stable synchronous states (i.e., order) and chaotic asynchronous states (i.e., disorder) receive benefits including high search efficiency, large dynamic range, high sensitivity and high flexibility. The high search efficiency in an uninformed condition [9] is a result of the large variance in step length distributions (Fig. 1F), i.e., the generation of short to long step lengths, which can be classified as an advantage in large repertoires of behavior [26]. The large dynamic range, high sensitivity, and adaptive switching come from the characteristics of instability and robustness. Instability requires the system to have a positive Lyapunov exponent, whereas robustness requires a small Lyapunov exponent. A small, positive Lyapunov exponent therefore produces both the ability to change and the high diffusiveness derived from a long step length (Fig. S4). Differences amplified by high diffusiveness consequently lead to different trajectories and positions in response to stimuli (Fig. 3A). High instability leads to masking small stimuli, whereas high robustness results in insensitivity to stimuli. It is for this reason that, although such properties have been reported in high-dimensional systems [27, 26], they are also observed in low-dimensional systems, such as the present one. Switching behavior can be related to cognitive functions. It has been shown that the critical state in brain dynamics is related to high cognitive abilities [66]. Being at a critical point should therefore be a fundamental part of creating adaptive behavior. From these results and from the literature on the critical point hypothesis, the possibility emerges that the main evolutionary force of Lévy walks is not search efficiency. It is worth noting that systems near a critical point receive some kinds of benefits simultaneously, and thus Lévy walks may be widely observed in nature.

Because we have proposed a model for Lévy walks, it is important to strictly identify the core network that plays a role in the Lévy walk generator in the brain and the parameters (e.g., neuromodulators) that control movement patterns [67, 30, 20]. Furthermore, although we have focused mainly on spontaneous behavior and behavioral change caused by stimuli, it is crucial to reveal how spontaneous dynamics, sensory inputs, and motor outputs are coupled with each other to be able to comprehensively understand the behavior of biological autonomous agents [68].

## Methods

### Model

To model Lévy walks based on nonlinear dynamics, we consider an autonomous agent with internal states that are composed of a minimal neural network-like system. The system has only two elements denoted by *x_t_*, *y_t_* ∈ [0, 1] and follows nonlinear dynamics with discrete time. The movement of the agent is determined by the internal states (see details below). We then observe the positions ***X***(*t*) = (*X_t_, Y_t_*) of the agents in a continuous two-dimensional space, and evaluate their movement trajectory.

We assume the update rules of the internal states to be the following equations:

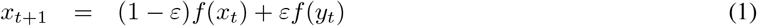

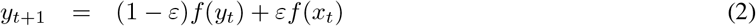

where *ε* ∈ [0.0, 0.5] is the coupling strength between two components *x* and *y*, and a control parameter of the system in our analysis. For *ε* = 0, *x_t_* and *y_t_* are independent of each other, whereas for *ε* = 0.5, *x_t_* and *y_t_* are the same. Here, we used a tent map as a nonlinear function *f* in each component because the tent map exhibits chaotic dynamics that can result in the randomness necessary for the random walks we seek here [22]. In addition, the tent map is tractable mathematically (the extension for the other form of the function *f* or a larger number of components is described in the Supplementary Information). We used the following tent map *f*:

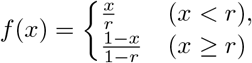

where *r* is a parameter that characterizes the deformation of tent maps. In our study, we set *r* = 0.7, but the main result does not qualitatively depend on *r* (Fig. S5). The general model for such systems includes globally coupled models that are used for modeling a class of nonlinear systems, including neural networks [43].

Finally, we simply define the movement of the agent determined by *x_t_* and *y_t_*, and assume the absolute angle *θ_t_* and the turning angle Δ*θ_t_*, of the agent at *t* are as follows:

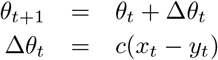

where *c* is a magnitude parameter that regulates the scale of the direction changes. We set *c* equal to *π* / max(|*x_t_*, – *y_t_*|) because the maximum difference between *x_t_* and *y_t_* should correspond to the largest turning angle, *π* or – *π*. Practically, *c* was obtained for each *ε* using more than 10^3^ time steps in advance. If max (|*x_t_*, – *y_t_*|) equaled zero (i.e., *ε* was in a stable region) we set *c* = *π*. This scenario is similar to the senario when the the maximum difference exceeds zero.

The speed of the movements of the agents is kept constant, and thus we assume that the speed is equal to one unit space per time step. The position ***X***(*t*) = (*X_t_, Y_t_*) of the agent can therefore be updated at each time *t* as follows:

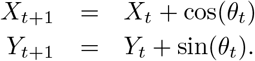

The larger the differences between the two outputs, the larger the turning angles. In contrast, consecutive small differences (e.g., approx. zero) between the outputs can produce movements of straight lines such that *θ*_*t*+1_ ≈ *θ_t_*. Note that we do not assume any specific probability distribution, including a power-law distribution, and thus the process is completely deterministic. Moreover, the parameter and function form of the system is symmetric for *x* and *y* because we expect unbiased movements to the right and left.

The initial conditions of *x* and *y* were drawn from a uniform distribution (0, 1) randomly and independently. We then ran and abandoned the first 10^3^ steps because of the possibility that the initial state was not on an attractor. We denoted the time after the first 10^3^ steps as *t* = 0. The initial position ***X***(0) = (*X*_0_, *Y*_0_) in the two-dimensional space was set to (0, 0). Because we considered the environmental space to be featureless with no borders, the agents obtained no stimuli from the environment. Even under such conditions, they could move around in space (Fig. 1B-D). Their movement was therefore classified as spontaneous behavior resulting from internal states such as brain dynamics.

### Evaluation of movement trajectory

To evaluate the movement trajectories produced by the agents, we used a rigorous method for fitting probability distributions to the step length distribution as well as empirical studies [34, 35, 36]. Humphries et al. [36] proposed a robust method for the analysis of movement trajectories in which the two-dimensional trajectory is mapped onto two one-dimensional axes, X and Y. We analyzed only one of these axes because our model is unbiased in terms of the X and Y axes. The step lengths of the consecutive and same (i.e., no turns) directional movements were then evaluated based on a maximum likelihood estimation (see the details in [35]). Note that the minimum step length was set to 1 in the analysis because we knew that it was 1 in the simulation.

Moreover, to characterize the property of diffusion, we calculated the mean squared displacement (MSD) defined as follows:

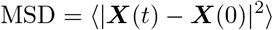

where 〈·〉 represents the ensemble average for trajectories with different initial states.

### Evaluation of dynamic ranges

To analyze the dynamic ranges, we added a stimulus with an amplitude *S* to the internal dynamics at *t_p_* in the form Δ*x* = *S* cos *ω* and Δ*y* = *S* sin *ω*, where *ω* is a random number drawn from a uniform distribution [−*π*, *π*]. Note that if *x* or *y* fell out of the domain [0, 1], the random number was generated again. We calculated the response *F* to *S* as the difference between the two positions of ***X***_O_(*t_p_* + *τ*) and ***X***_P_(*t_p_* + *τ*) (Fig. 3A). Then, to evaluate how to respond to the stimulus, the dynamic range was defined as Δ*_S_* = 10 log_10_(*S*_0.9_/*S*_0.1_) where *S_z_* is the size of a perturbation that produces *F_z_* = *F*_min_ + *z*(*F*_max_ – *F*_min_) (see the inset of Fig. 3C).

For switching behavior, we started the dynamics from an initial states *x_t_p__* = 0.5 + *S*′ sin *ω* and *y_t_p__* = 0.5 + *S*′ cos *ω*, where *S′* is a parameter controlling the types of stimuli, and *ω* is a uniform random number [−*π*, *π*]. A small *S′* generates a straight movement because Δ*θ*(∝ *x* – *y*) is close to 0. In contrast, a large *S′* can generate a turn. The distance *G* between ***X***(*t_p_*) and ***X***(*t_p_* + *τ*) was calculated to quantify the degree of exploitation/exploration (Fig. 4A). In addition to Δ*_S_*, we defined the dynamic range for *S′* as 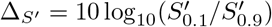 where 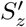 is a stimulus parameter that produces *G_z_ G*_min_ + *z*(*G*_max_ – *G*_min_).

### Analysis on empirical data

To verify the result from the proposed model, we analyzed time series of turning angles in the movement trajectories from a published dataset of freely moving *Drosophila* larvae [20, 69]. Our analysis included all conditions such as with/without blocked neurons and temperatures. To compare with the results of the constant speed model, we transformed the raw data and redefined the direction change angles for each constant step length. We used three times the mean movement distance per second as constant step length. For other values, the main results did not change qualitatively. In addition, movements with unusually fast speed (here more than three times the mean speed), which is considered a tracking error, were excluded from the time series. Finally, we removed time series with fewer than 100 data points and used 291 of 304 time series to investigate nonlinearity.

We analyzed time series by using a state space reconstruction method based on Takens’ embedding theorem [46]. In the analysis, the original time series *z_i_*(*i* = 1, …, *N*), where *N* is the length of time series, was transformed into lagged coordinate vectors ***v***(*i*) = {*z_i_*, *z_i–T_*, …, *z*_*i*–(*E*–1)*T*_} where *E* is an embedding dimension and *T* is a time lag (here we used *T* = 1). To obtain the embedding dimension of time series that best estimates dimension of attractor manifold, we used a simplex projection (the range of E was 1 to 10) [47]. We then determined whether the time series was generated from nonlinear dynamics or stochastic processes by using S-map with the best embedding dimension [47]. S-map is a prediction method with a parameter *ϕ*(≥ 0) that controls the nonlinearity of the prediction model. For *ϕ* = 0, the prediction model produces linear predictions using data points with equal weights. In contrast, the model for *ϕ* > 0 uses locally weighted (i.e., state dependent) data points for prediction. Thus, if the model with *ϕ* > 0 produces higher prediction accuracy (correlation coefficient *ρ* between observed and predicted time series), we can conclude that the time series is derived from nonlinear dynamics. In addition, we conducted a statistical test using phase-randomized (i.e., linear) surrogate time series to investigate significant increases of Δ*ρ*_max_ = *ρ*_*ϕ*=*ϕ*\*_ – *ρ*_*ϕ*=0_, where *ϕ*\* is a value that produces the maximum *ρ*. Then, we calculated *p*-values by identifying the position of empirical Δ*ρ*_max_ among surrogate data (100 replicates). Once we confirmed that a time series was generated from nonlinear dynamics, we calculated the largest Lyapunov exponents by using Kantz algorithm (Fig. S9A) [70]. In the analysis, we used the embedding dimension estimated by simplex projection. Likewise, Kantz algorithm was used for the time series from the proposed model to compare the Lyapunov exponents.

Finally, to reveal responses to perturbations in real organisms, we developed a data-based method based on the idea of the twin surrogate method [48]. This makes it possible to simulate the changes of time series by perturbations only from observational data. After we constructed lagged coordinated vectors from empirical time series by using the best *E*, we looked for indistinguishable points in reconstructed phase space. These points are called twins (see the details in Supplementary Information). By randomly switching the trajectory to another trajectory at the point with a twin pair, we could generate another time series from the same system (i.e., attractor) with the original time series (see the details in Supplementary Information). This method is an original twin surrogate method [48].

To examine *F* for the small *S* case, we started with a randomly chosen point, and then we generated a subsequent time series with the twin surrogate method. Based on the generated time series, a movement trajectory was created and compared to the position of the original movement trajectory with the same initial point after a time *τ*. In the case of a large perturbation (corresponding to the large *S* case), the starting point was randomly selected from the points of the embedded vectors ***v*** that fell within the top 5% of the distribution of ‖***v***‖ where ‖ · ‖ means the Euclidean norm, and subsequent time series were generated with the twin surrogate method and similarly compared to the original movement trajectory. To examine behavioral switching according to the type of stimulus, we randomly selected initial values from the bottom 5% and top 5% of the distribution of ‖ ***v*** ‖ to correspond to the small and large *S*′ case, respectively (Fig. 4). We then generated subsequent time series with the twin surrogate method in order to calculate the distance between the initial position and the position after *τ*. In these analyses, we used 100 replications for each original time series. See the details of this method in Supplementary Information.

## Data Availability

All data analysed in this study are available in Dryad (http://doi.org/10.5061/dryad.7m0cfxpq0) [69].

## Notes

### Competing Interest Statement

The authors have declared no competing interest.

## References

[1] Gandhimohan M Viswanathan, Marcos GE Da Luz, Ernesto P Raposo, and H Eugene Stanley. The physics of foraging: an introduction to random searches and biological encounters. Cambridge University Press, 2011.

[2] Tajie H Harris, Edward J Banigan, David A Christian, Christoph Konradt, Elia D Tait Wojno, Kazumi Norose, Emma H Wilson, Beena John, Wolfgang Weninger, Andrew D Luster, et al. Generalized lévy walks and the role of chemokines in migration of effector cd8+ t cells. Nature, 486(7404):545, 2012.

[3] Sabil Huda, Bettina Weigelin, Katarina Wolf, Konstantin V Tretiakov, Konstantin Polev, Gary Wilk, Masatomo Iwasa, Fateme S Emami, Jakub W Narojczyk, Michal Banaszak, et al. Lévy-like movement patterns of metastatic cancer cells revealed in microfabricated systems and implicated in vivo. Nature communications, 9(1):4539, 2018.

[4] Nicolas E Humphries, Nuno Queiroz, Jennifer RM Dyer, Nicolas G Pade, Michael K Musyl, Kurt M Schaefer, Daniel W Fuller, Juerg M Brunnschweiler, Thomas K Doyle, Jonathan DR Houghton, et al. Environmental context explains lévy and brownian movement patterns of marine predators. Nature, 465(7301):1066, 2010.

[5] David A Raichlen, Brian M Wood, Adam D Gordon, Audax ZP Mabulla, Frank W Marlowe, and Herman Pontzer. Evidence of lévy walk foraging patterns in human hunter–gatherers. Proceedings of the National Academy of Sciences, 111(2):728–733, 2014.

[6] Andy Reynolds, Eliane Ceccon, Cristina Baldauf, Tassia Karina Medeiros, and Octavio Miramontes. Lévy foraging patterns of rural humans. PLOS ONE, 13(6):e0199099, 2018.

[7] Christopher T Kello, Gordon DA Brown, Ramon Ferrer-i Cancho, John G Holden, Klaus Linkenkaer-Hansen, Theo Rhodes, and Guy C Van Orden. Scaling laws in cognitive sciences. Trends in cognitive sciences, 14(5):223–232, 2010.

[8] Theo Rhodes and Michael T Turvey. Human memory retrieval as lévy foraging. Physica A: Statistical Mechanics and its Applications, 385(1):255–260, 2007.

[9] Gandimohan M Viswanathan, Sergey V Buldyrev, Shlomo Havlin, MGE Da Luz, EP Raposo, and H Eugene Stanley. Optimizing the success of random searches. Nature, 401(6756):911, 1999.

[10] Frederic Bartumeus, M G E da Luz, Gandhimohan M Viswanathan, and Jordi Catalan. Animal search strategies: a quantitative random-walk analysis. Ecology, 86(11):3078–3087, 2005.

[11] ME Wosniack, MC Santos, EP Raposo, GM Viswanathan, and MGE Da Luz. Robustness of optimal random searches in fragmented environments. Physical Review E, 91(5):052119, 2015.

[12] Marina E Wosniack, Marcos C Santos, Ernesto P Raposo, Gandhi M Viswanathan, and Marcos GE da Luz. The evolutionary origins of lévy walk foraging. PLOS Comput. Biol., 13(10):e1005774, 2017.

[13] Andy Reynolds. Liberating lévy walk research from the shackles of optimal foraging. Physics of life reviews, 14:59–83, 2015.

[14] Andy M Reynolds. Current status and future directions of lévy walk research. Biology open, 7(1):bio030106, 2018.

[15] Alexander Maye, Chih-hao Hsieh, George Sugihara, and Björn Brembs. Order in spontaneous behavior. PLOS ONE, 2(5):e443, 2007.

[16] Alex Proekt, Jayanth R Banavar, Amos Maritan, and Donald W Pfaff. Scale invariance in the dynamics of spontaneous behavior. Proceedings of the National Academy of Sciences, 109(26):10564–10569, 2012.

[17] Victoria J Wearmouth, Matthew J McHugh, Nicolas E Humphries, Aurore Naegelen, Mohammed Z Ahmed, Emily J Southall, Andrew M Reynolds, and David W Sims. Scaling laws of ambush predator ‘waiting’behaviour are tuned to a common ecology. Proceedings of the Royal Society B: Biological Sciences, 281(1782):20132997, 2014.

[18] Andrea Kölzsch, Adriana Alzate, Frederic Bartumeus, Monique de Jager, Ellen J Weerman, Geerten M Hengeveld, Marc Naguib, Bart A Nolet, and Johan van de Koppel. Experimental evidence for inherent lévy search behaviour in foraging animals. Proceedings of the Royal Society B: Biological Sciences, 282(1807):20150424, 2015.

[19] Naohisa Nagaya, Nobuaki Mizumoto, Masato S Abe, Shigeto Dobata, Ryota Sato, and Ryusuke Fujisawa. Anomalous diffusion on the servosphere: A potential tool for detecting inherent organismal movement patterns. PLOS ONE, 12(6):e0177480, 2017.

[20] David W Sims, Nicolas E Humphries, Nan Hu, Violeta Medan, and Jimena Berni. Optimal searching behaviour generated intrinsically by the central pattern generator for locomotion. eLife, 8:e50316, 2019.

[21] Blaine J Cole. Is animal behaviour chaotic? evidence from the activity of ants. Proceedings of the Royal Society of London. Series B: Biological Sciences, 244(1311):253–259, 1991.

[22] Andy M Reynolds, Frederic Bartumeus, Andrea Kölzsch, and Johan Van De Koppel. Signatures of chaos in animal search patterns. Scientific reports, 6:23492, 2016.

[23] Esther D Gutiérrez and Juan Luis Cabrera. A neural coding scheme reproducing foraging trajectories. Scientific reports, 5:18009, 2015.

[24] Theo Geisel and Stefan Thomae. Anomalous diffusion in intermittent chaotic systems. Physical review letters, 52(22):1936, 1984.

[25] Michael F Shlesinger, George M Zaslavsky, and Joseph Klafter. Strange kinetics. Nature, 363(6424):31–37, 1993.

[26] Miguel A Munoz. Colloquium: Criticality and dynamical scaling in living systems. Reviews of Modern Physics, 90(3):031001, 2018.

[27] Osame Kinouchi and Mauro Copelli. Optimal dynamical range of excitable networks at criticality. Nature physics, 2(5):348–351, 2006.

[28] Auke Jan Ijspeert. Central pattern generators for locomotion control in animals and robots: a review. Neural networks, 21(4):642–653, 2008.

[29] Eve Marder, Dirk Bucher, David J Schulz, and Adam L Taylor. Invertebrate central pattern generation moves along. Current Biology, 15(17):R685–R699, 2005.

[30] Jimena Berni. Genetic dissection of a regionally differentiated network for exploratory behavior in drosophila larvae. Current Biology, 25(10):1319–1326, 2015.

[31] Valentin A Nepomnyashchikh and Konstantin A Podgornyj. Emergence of adaptive searching rules from the dynamics of a simple nonlinear system. Adaptive Behavior, 11(4):245–265, 2003.

[32] Hatsuo Hayashi, Satoru Ishizuka, Masahiro Ohta, and Kazuyoshi Hirakawa. Chaotic behavior in the onchidium giant neuron under sinusoidal stimulation. Physics Letters A, 88(8):435–438, 1982.

[33] George J Mpitsos, Robert M Burton Jr, H Clayton Creech, and Seppo O Soinila. Evidence for chaos in spike trains of neurons that generate rhythmic motor patterns. Brain Research Bulletin, 21(3):529–538, 1988.

[34] Aaron Clauset, Cosma Rohilla Shalizi, and Mark EJ Newman. Power-law distributions in empirical data. SIAM review, 51(4):661–703, 2009.

[35] Nicolas E Humphries, Henri Weimerskirch, Nuno Queiroz, Emily J Southall, and David W Sims. Foraging success of biological lévy flights recorded in situ. Proceedings of the National Academy of Sciences, 109(19):7169–7174, 2012.

[36] Nicolas E Humphries, Henri Weimerskirch, and David W Sims. A new approach for objective identification of turns and steps in organism movement data relevant to random walk modelling. Methods in Ecology and Evolution, 4(10):930–938, 2013.

[37] Frederic Bartumeus and Simon A Levin. Fractal reorientation clocks: Linking animal behavior to statistical patterns of search. Proceedings of the National Academy of Sciences, 105(49):19072–19077, 2008.

[38] Arkady S Pikovsky and Peter Grassberger. Symmetry breaking bifurcation for coupled chaotic attractors. Journal of Physics A: Mathematical and General, 24(19):4587, 1991.

[39] Edward Ott and John C Sommerer. Blowout bifurcations: the occurrence of riddled basins and on-off intermittency. Physics Letters A, 188(1):39–47, 1994.

[40] Arkady Pikovsky, Jurgen Kurths, Michael Rosenblum, and Jürgen Kurths. Synchronization: a universal concept in nonlinear sciences, volume 12. Cambridge university press, 2003.

[41] Björn Brembs. Towards a scientific concept of free will as a biological trait: spontaneous actions and decisionmaking in invertebrates. Proceedings of the Royal Society B: Biological Sciences, 278(1707):930–939, 2010.

[42] Robert M May. Simple mathematical models with very complicated dynamics. Nature, 261(5560):459, 1976.

[43] Kunihiko Kaneko. Clustering, coding, switching, hierarchical ordering, and control in a network of chaotic elements. Physica D: Nonlinear Phenomena, 41(2):137–172, 1990.

[44] Thomas T Hills, Peter M Todd, David Lazer, A David Redish, Iain D Couzin, Cognitive Search Research Group, et al. Exploration versus exploitation in space, mind, and society. Trends in cognitive sciences, 19(1):46–54, 2015.

[45] Simon Benhamou. How many animals really do the lévy walk? Ecology, 88(8):1962–1969, 2007.

[46] Floris Takens. Detecting strange attractors in turbulence. In Dynamical systems and turbulence, Warwick 1980, pages 366–381. Springer, 1981.

[47] Chih-hao Hsieh, Sarah M Glaser, Andrew J Lucas, and George Sugihara. Distinguishing random environmental fluctuations from ecological catastrophes for the north pacific ocean. Nature, 435(7040):336–340, 2005.

[48] Marco Thiel, Maria Carmen Romano, Jürgen Kurths, Martin Rolfs, and Reinhold Kliegl. Twin surrogates to test for complex synchronisation. EPL (Europhysics Letters), 75(4):535, 2006.

[49] Vijay Mohan K Namboodiri, Joshua M Levy, Stefan Mihalas, David W Sims, and Marshall G Hussain Shuler. Rationalizing spatial exploration patterns of wild animals and humans through a temporal discounting framework. Proceedings of the National Academy of Sciences, 113(31):8747–8752, 2016.

[50] Łukasz Kuśmierz and Taro Toyoizumi. Emergence of lévy walks from second-order stochastic optimization. Physical review letters, 119(25):250601, 2017.

[51] Gyorgy Buzsaki. Rhythms of the Brain. Oxford University Press, 2006.

[52] Per Bak, Chao Tang, and Kurt Wiesenfeld. Self-organized criticality: An explanation of the 1/f noise. Physical review letters, 59(4):381, 1987.

[53] CL Faustino, LR Da Silva, MGE Da Luz, EP Raposo, and GM Viswanathan. Search dynamics at the edge of extinction: anomalous diffusion as a critical survival state. EPL (Europhysics Letters), 77(3):30002, 2007.

[54] Frederic Bartumeus, Jordi Catalan, UL Fulco, ML Lyra, and GM Viswanathan. Optimizing the encounter rate in biological interactions: Lévy versus brownian strategies. Physical Review Letters, 88(9):097901, 2002.

[55] Vladimir V Palyulin, Aleksei V Chechkin, and Ralf Metzler. Lévy flights do not always optimize random blind search for sparse targets. Proceedings of the National Academy of Sciences, 111(8):2931–2936, 2014.

[56] Masato S Abe and Masakazu Shimada. Lévy walks suboptimal under predation risk. PLOS Comput. Biol., 11(11):e1004601, 2015.

[57] Nobuaki Mizumoto, Masato S Abe, and Shigeto Dobata. Optimizing mating encounters by sexually dimorphic movements. Journal of The Royal Society Interface, 14(130):20170086, 2017.

[58] Nicolas Levernier, Johannes Textor, Olivier Bénichou, and Raphaël Voituriez. Inverse square lévy walks are not optimal search strategies for d ≥ 2. Physical Review Letters, 124(8):080601, 2020.

[59] Teodoro Dannemann, Denis Boyer, and Octavio Miramontes. Lévy flight movements prevent extinctions and maximize population abundances in fragile lotka–volterra systems. Proceedings of the National Academy of Sciences, 115(15):3794–3799, 2018.

[60] Thierry Mora and William Bialek. Are biological systems poised at criticality? Journal of Statistical Physics, 144(2):268–302, 2011.

[61] Jorge Hidalgo, Jacopo Grilli, Samir Suweis, Miguel A Munoz, Jayanth R Banavar, and Amos Maritan. Information-based fitness and the emergence of criticality in living systems. Proceedings of the National Academy of Sciences, 111(28):10095–10100, 2014.

[62] Bryan C Daniels, David C Krakauer, and Jessica C Flack. Control of finite critical behaviour in a small-scale social system. Nature communications, 8:14301, 2017.

[63] Bryan C Daniels, Hyunju Kim, Douglas Moore, Siyu Zhou, Harrison B Smith, Bradley Karas, Stuart A Kauffman, and Sara I Walker. Criticality distinguishes the ensemble of biological regulatory networks. Physical review letters, 121(13):138102, 2018.

[64] Ofer Feinerman, Itai Pinkoviezky, Aviram Gelblum, Ehud Fonio, and Nir S Gov. The physics of cooperative transport in groups of ants. Nature Physics, 14(7):683, 2018.

[65] Joschka Boedecker, Oliver Obst, Joseph T Lizier, N Michael Mayer, and Minoru Asada. Information processing in echo state networks at the edge of chaos. Theory in Biosciences, 131(3):205–213, 2012.

[66] Takahiro Ezaki, Elohim Fonseca dos Reis, Takamitsu Watanabe, Michiko Sakaki, and Naoki Masuda. Closer to critical resting-state neural dynamics in individuals with higher fluid intelligence. Communications biology, 3(1):1–9, 2020.

[67] Jean-Rene Martin, Philippe Faure, and Roman Ernst. The power law distribution for walking-time intervals correlates with the ellipsoid-body in drosophila. Journal of neurogenetics, 15(3–4):205–219, 2001.

[68] Karl Friston. Life as we know it. Journal of the Royal Society Interface, 10(86):20130475, 2013.

[69] Jimena Berni, David W Sims, and Nicolas E Humphries. Optimal searching behaviour generated intrinsically by the central pattern generator for locomotion. Dryad, v7, 2020.

[70] Rainer Hegger, Holger Kantz, and Thomas Schreiber. Practical implementation of nonlinear time series methods: The tisean package. Chaos: An Interdisciplinary Journal of Nonlinear Science, 9(2):413–435, 1999.

