## Supplementary information for "Functional advantages of Lévy walks emerging near a critical point"

### Supporting Information Text

**Mean squared displacement.** To characterize the diffusiveness of movements, we calculated the mean squared displacement (MSD) as defined in the Materials and Methods section of the main text for the simulated trajectories. Moreover, how MSD increases with time $t$ can characterize the movements (1) as follows:

$$\mathrm{MSD} = \langle (\boldsymbol{X_t} - \boldsymbol{X_0})^2 \rangle \sim t^{\alpha}$$

where $\alpha \in [0, 2]$ is an exponent for MSD. Brownian motions and straight lines have $\alpha = 1$ and $\alpha = 2$, respectively. The fact that the exponent $\alpha$ for Lévy walks is larger than 1 suggests that Lévy walks are superdiffusive.

For $\varepsilon = 0.1, 0.22, 0.3$, an ensemble of 100 trajectories was obtained from different initial conditions, $x_0$ and $y_0$, which were randomly chosen. The result shown in Fig. S1 shows that MSD and indicates that the slope $\alpha$ of $\varepsilon = 0.22$ is between those corresponding to $\varepsilon = 0.1$ and $\varepsilon = 0.3$. The relationship between time and MSD for $\varepsilon = 0.22$ does not seem linear over the whole range. The reason is that the step length distribution is not a pure power law distribution; instead, it a truncated power law distribution. Therefore, the slope of the log(MSD) versus log(time) relationship can converge to 1 after a sufficiently long time. However, we can conclude that the movement patterns for $\varepsilon = 0.22$ are highly diffusive; they are more diffuse than those for $\varepsilon = 0.1$, and they are less diffuse than those for $\varepsilon = 0.3$, considering that biologically realistic situations are not the limit of long time. This pattern is consistent with the argument that the movement patterns can be classified as Lévy walks.

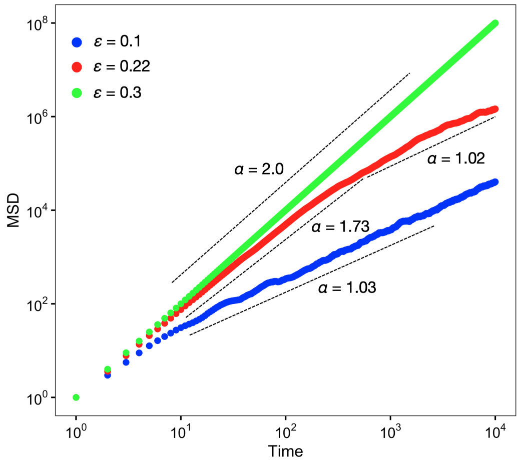


**Fig. S1.** Mean squared displacement (MSD). Blue dots represent MSD for $\varepsilon = 0.1$, red dots represent MSD for $\varepsilon = 0.22$, and green dots represent MSD for $\varepsilon = 0.3$ The dashed lines represent the regression lines with the slope $\alpha$.

**Characteristics of dynamics.** We showed that the movement trajectories of our model (Eqs. 1 and 2 in the main text) can be classified into Lévy walks, Brownian motions, and ballistic movements (see main text). To intuitively grasp how the dynamics of the internal states $x_t, y_t$ produce such variable movement patterns, we show the attractors of $x_t, y_t$ (Fig. S2). The diagonal line in Fig. S2 represents $x = y$, namely, synchronous states that lead to no turnings in the movements. Thus, if $x = y$, namely, $\Delta\theta = 0$ is a (globally) stable attractor, the movement patterns are equivalent to straight (ballistic) movements ($\varepsilon = 0.3$; Fig. S2C), although $x$ and $y$ fluctuate in a chaotic manner. In contrast, if $x = y$ is unstable, $x - y$ exhibits random fluctuations (see insets of Fig. 1BC in the main text). In particular, the case of $\varepsilon = 0.1$ covers a large range of $x$ and $y$ (Fig. S2A). Consequently, the movement patterns become a normal diffusion process, that is, Brownian motions. Between these two patterns ($\varepsilon = 0.22$), we can see the mixture of $x \simeq y$ and random fluctuations, which produces intermittent patterns in the dynamics of $x - y$ and Lévy walks as movement trajectories (Fig. S2B).

**Stability condition for ballistic movements.** To understand how the phase transition shown in Fig. 2 of the main text occurs, we derive the stability condition for ballistic movements, that is, $\Delta\theta = 0$. First, we set the new variables to $H = (x + y)/2$ and $K = (x - y)/2$. From Eqs. 1 and 2 in the main text, we obtain the difference equation for the new variables as follows:

$$H(t+1) = \frac{1}{2}(f(H(t) + K(t)) + f(H(t) - K(t)))$$
$$K(t+1) = \frac{1-2\varepsilon}{2}(f(H(t) + K(t)) - f(H(t) - K(t)))$$

where $f$ is a nonlinear function that produces chaotic dynamics. Here, as mentioned in the main text, we used a tent map. The equations of a small perturbation $h, k$ in the vicinity of $G = 0$ are obtained by linearization as follows:

$$h(t+1) = f'(H(t))h(t)$$
$$k(t+1) = (1 - 2\varepsilon)f'(H(t))k(t)$$

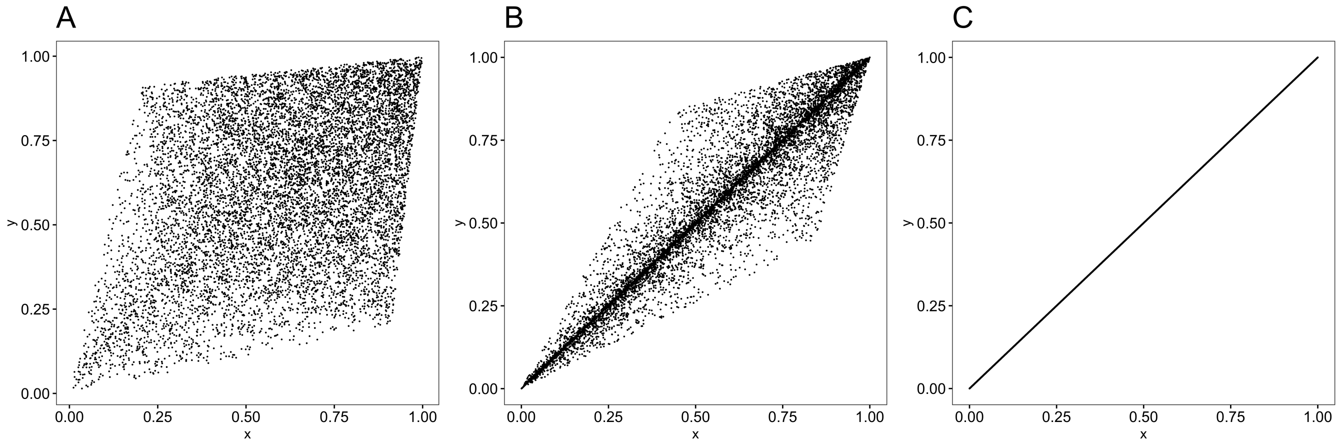


**Fig. S2.** Attractors depending on the parameter $\varepsilon$. The horizontal and vertical axes represent the internal states $x$ and $y$ of the agent, respectively. The black dots represent the value at a certain time. (A) $\varepsilon = 0.1$, that is, attractor of Brownian motions. (B) $\varepsilon = 0.22$, that is, attractor of Lévy walks. (C) $\varepsilon = 0.3$, attractor of straight movement. The other parameters were set to $r = 0.7$ and $t_{\max} = 10{,}000$.

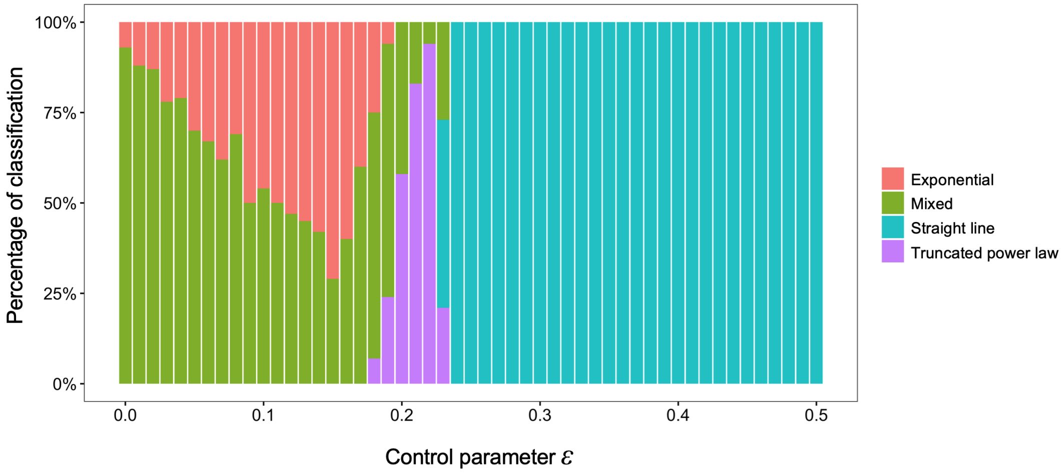


**Fig. S3.** Detail of the classifications based on statistical analysis. The data are the same as shown in Fig. 2 in the main text.

Note that we can separate $k(t)$ from $h(t)$. Because $H(t+1) = f(H(t))$ exhibits chaotic dynamics, the linear stability for $x - y = 0$ can be characterized by two exponents:

$$\lambda_h = \langle \ln |f'(H)| \rangle$$
$$\lambda_k = \ln |1 - 2\varepsilon| + \langle \ln |f'(H)| \rangle.$$

where $\varepsilon$ is the coupling strength in the system, and $\langle \ln |f'(H)| \rangle$ denotes a Lyapunov exponent of the nonlinear map $f$ because of the ergodicity of chaos (2). Because a variable in a non-coupled tent map $f$ is distributed uniformly from 0 to 1 (2), we can analytically derive the Lyapunov exponents $\lambda_{tent} = \lambda_h$ of a single tent map as follows:

$$\lambda_{tent} = -r \ln r - (1 - r) \ln (1 - r)$$

where $r$ is a parameter of the tent map. From the above, the stability index $\lambda_c$ of $x - y$ is described as

$$\lambda_c = \ln |1 - 2\varepsilon| + \lambda. \qquad [1]$$

Therefore, the stability condition for $\Delta\theta = c(x-y) = 0$ is $\ln |1 - 2\varepsilon| + \lambda < 0$. For the case of $r = 0.7$, we obtain $\varepsilon > \varepsilon_c \simeq 0.228$ as the stability condition, which is consistent with the main result.

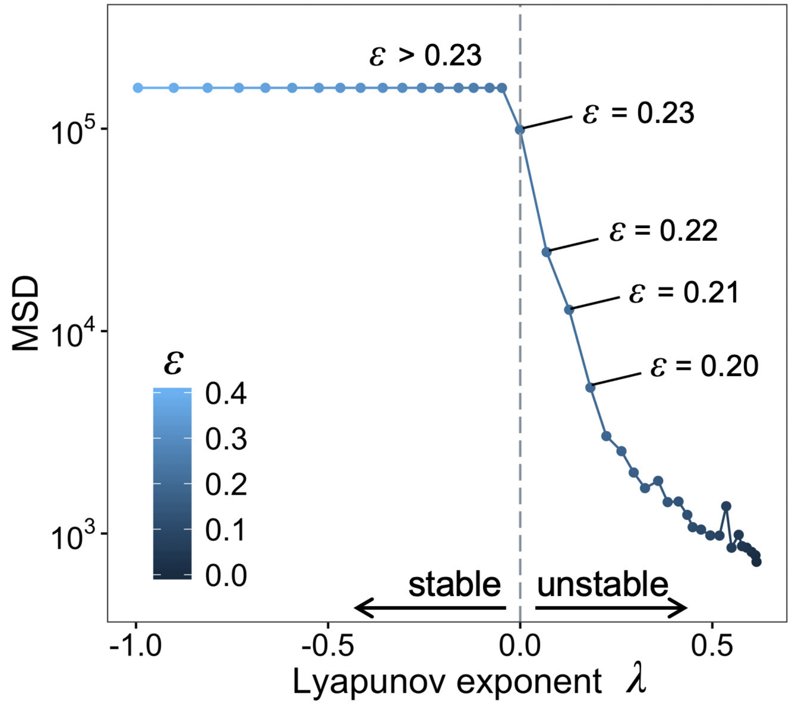


**Fig. S4.** A trade-off relationship between the instability index (Lyapunov exponent $\lambda$) of $x_t - y_t$ and the diffusiveness (MSD) of the agents. The Lyapunov exponent was obtained analytically using equation 1. The MSD was calculated for $t = 400$. The color of the plots represents the control parameter $\varepsilon$. Note that the results for $\varepsilon > 0.4$ are not shown to improve the visibility. The gray dashed line corresponds to a Lyapunov exponent of 0, which signifies a boundary between the stable and unstable phases.

**Other models.** We assessed the robustness of the results for emergence conditions of Lévy walks as shown in Fig. 2 of the main text for different nonlinear functions, the number of elements, and stochastic noise.

***Model with different nonlinear functions.*** First, we explored the effect of the parameter $r$ in the tent map on the condition for the emergence of Lévy walks. Fig. S5 shows that the parameter $r$ does not qualitatively affect the result. In the case of $r = 0.5$, we used $r = 0.501$ because the rounding error of the values caused a falling to a fixed point. The result suggests that the tendency of Lévy walks to emerge near a critical point was insensitive to $r$. Second, we used different nonlinear functions, here logistic maps $f(x) = ax(1 - x)$, where $a \in [0, 4]$ is a parameter, to determine the attractor (3). The result of the model with $a = 3.9$, which produces chaotic dynamics, is shown in Fig. S6 and also supports the main conclusion.

***Model with large degree of freedom.*** In the main analysis, we adopted a model system composed of only two elements. However, one might wonder if the number of elements affects the result. To investigate this question, we analyzed globally coupled maps

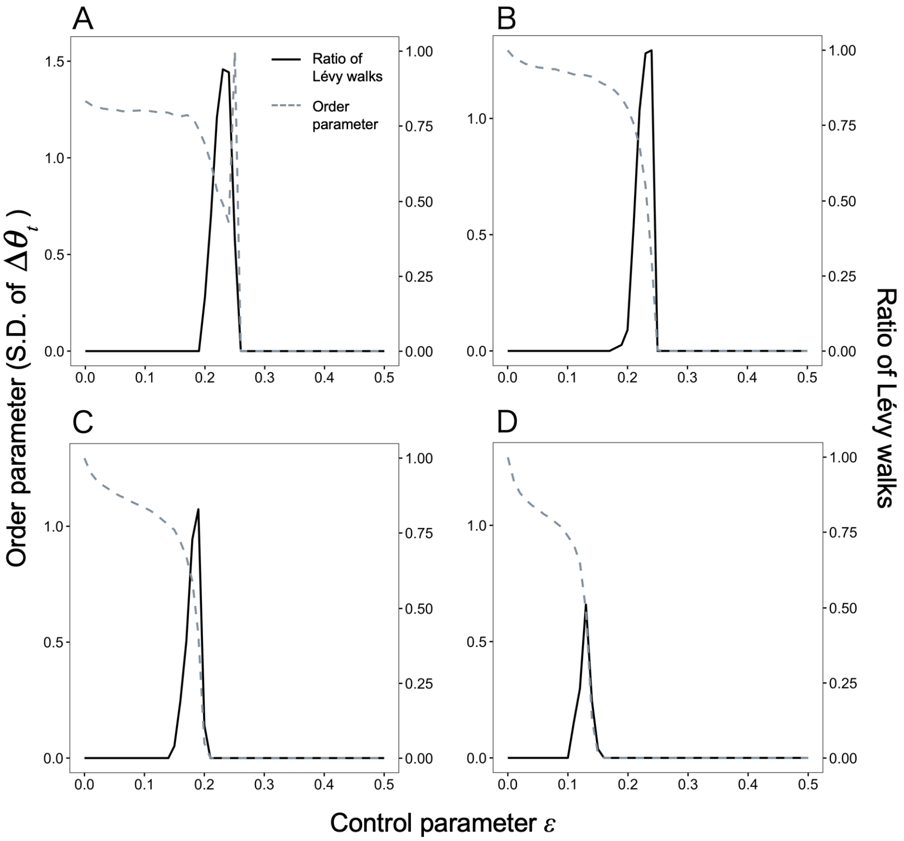


**Fig. S5.** Phase diagram for the model with different $r$. (A)$r = 0.501$, (B) $r = 0.6$, (C) $r = 0.8$, (D) $r = 0.9$. The other settings are the same as for the results shown in Fig. 2 in the main text.

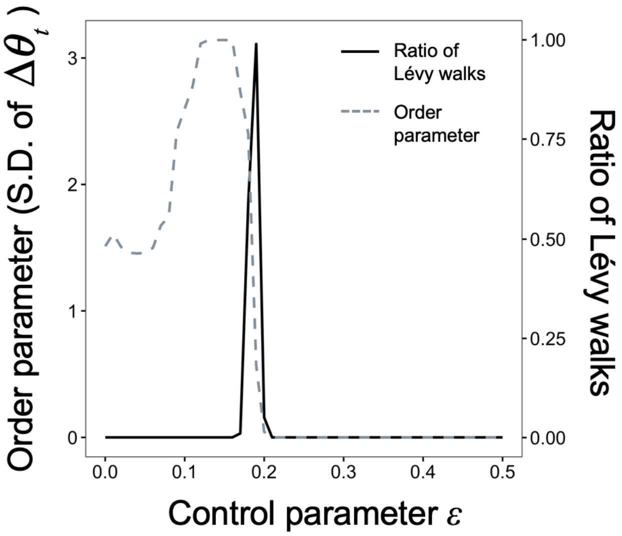


**Fig. S6.** Phase diagram for the logistic map model with $a = 3.9$. The other settings are the same as those shown in the results in Fig. 2 in the main text.

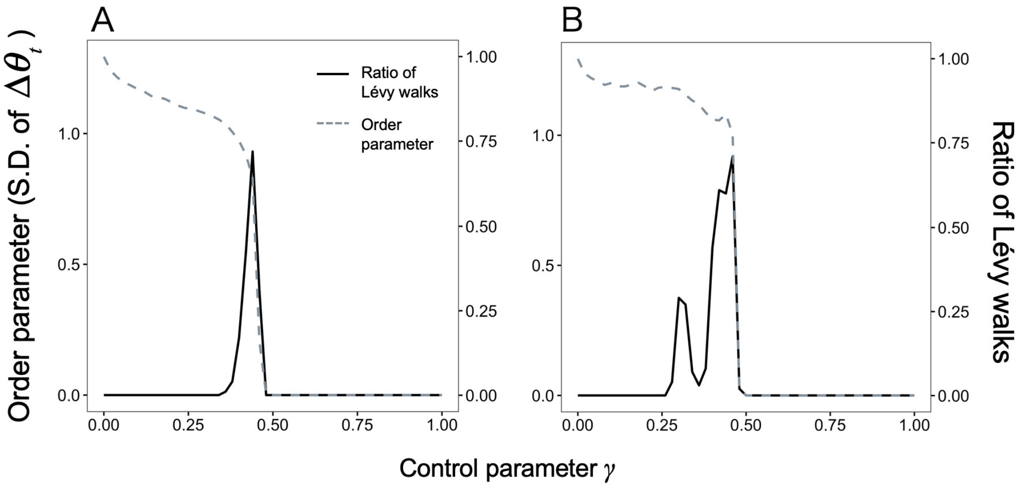


**Fig. S7.** Phase diagram for the model with large degrees of freedom. (A) $N = 10$, (B) $N = 100$. The other settings are the same as those shown in Fig. 2 in the main text.

(GCMs) with $N$ elements, which have been used for modeling complex systems such as neural networks (4). The definition of GCMs is as follows:

$$x_i(t+1) = (1-\gamma)f(x_i(t)) + \frac{\gamma}{N}\sum_{j=1}^{N} f(x_j(t))$$

where $x_i$ is a value of element $i$ and $\gamma \in [0,1]$ is the coupling strength between the elements. Here, as we did in the main analysis, we used a tent map with $r = 0.7$ as the nonlinear function $f$. Note that our model in the main analysis was a special case for $N = 2$ when $\gamma = 2\varepsilon$. We randomly chose two elements as the output nodes and used them for producing the movements. We used the same procedure for the two-elements model (Fig. 1A in the main text). The results for $N = 10$ and 100 are shown in Fig. S7. The results suggest that even for systems with a relatively high number of elements, Lévy walks can emerge near the critical point.

***Model with noise.*** When one considers a biologically realistic situation, there is noise in the system. To verify the robustness of the result of the emergence of Lévy walks near the critical point, we added an inherently stochastic noise to the systems. At each time step, white noise following a normal distribution with $\mu = 0, \sigma = \eta$ was added to the internal states $x_t$ and $y_t$ independently. Fig. S8 shows that although the stable area for large $\varepsilon$ is likely to break down with increasing size of the noise, the tendency for the emergence of Lévy walks does not change qualitatively.

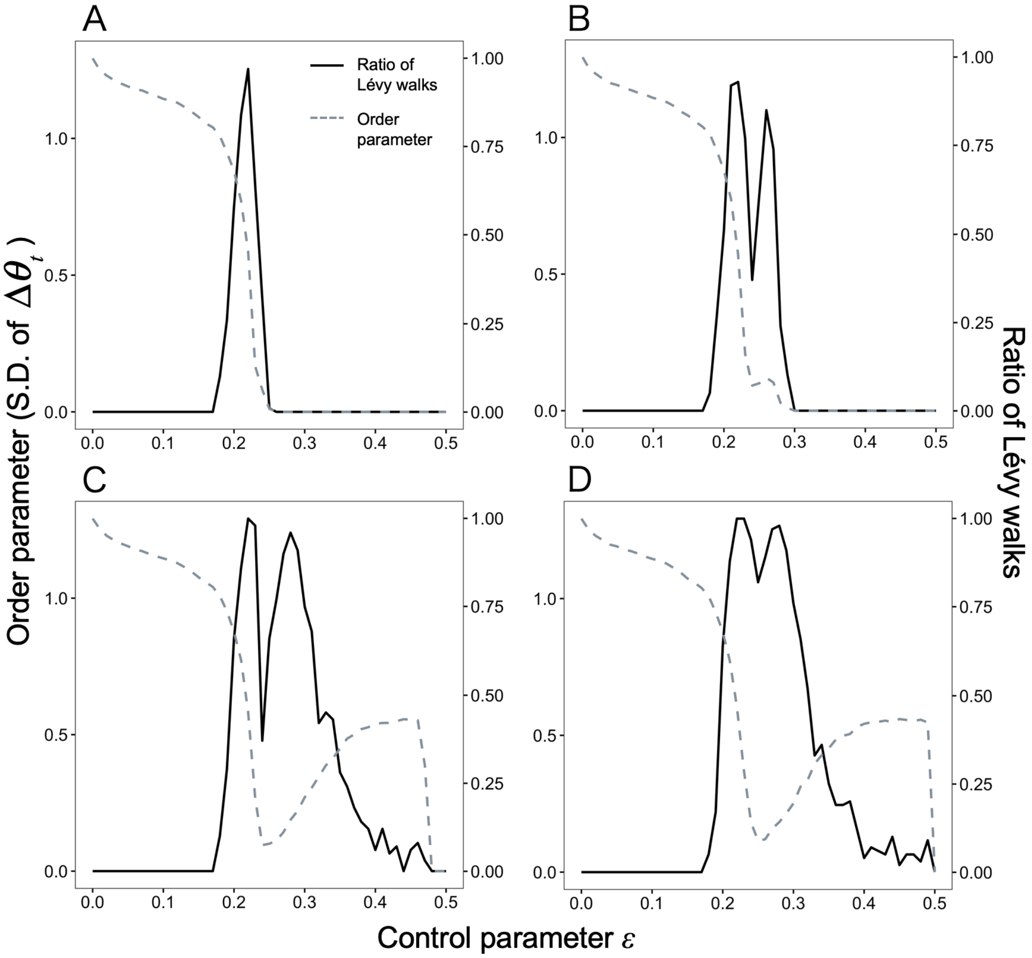


**Fig. S8.** Phase diagram for the model with stochastic noise. (A) $\eta = 1.0 \times 10^{-10}$, (B) $\eta = 1.0 \times 10^{-8}$, (C) $\eta = 1.0 \times 10^{-6}$, (D) $\eta = 1.0 \times 10^{-4}$. The other settings are the same as those shown in the results in Fig. 2 in the main text.

**Data-based simulation of responses to perturbations.** To examine responses to a perturbation in the empirical time series $z_1, \ldots, z_N$ of turning angles (Fig. 5B in the main text) where $N$ is the number of data points, we developed a method based on a twin surrogate algorithm proposed by Thiel et al. (5). The twin surrogate algorithm allowed us to produce surrogate data with different initial states while preserving the properties of a nonlinear dynamical system of the original time series. In addition to this algorithm, we virtually observed the response to a perturbation by specifying the initial value corresponding to

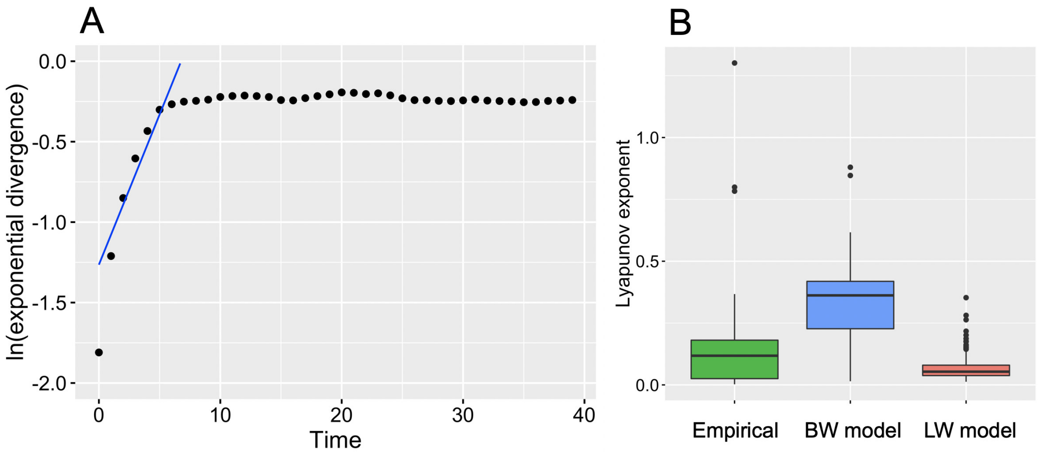


**Fig. S9.** Lyapunov exponents in empirical data. (A) The Lyapunov exponent is estimated as the initial slope of the log-transformed exponential divergence versus time, as calculated by linear regression (blue line) for a linear increasing region. In our analysis, a regression line was obtained for a value $d(\geq 1)$ up to 95% of the maximum value of the vertical axis. We changed $d$ and selected the regression line with $R^2$ values greater than 0.9 that fit the longest region. Note that data with the longest fitted region of 3 or less were excluded. (B) Estimated Lyapunov exponents in empirical data ($n = 158$), Brownian walk model (BW; $\varepsilon = 0.0, n = 246$), and Lévy walk model (LW; $\varepsilon = 0.22, n = 253$). The result of the multiple comparison test showed that all the differences between them were significant ($p < 10^{-5}$).

the perturbation size and stimuli type in the theoretical analysis (Figs. 3–4 in the main text) and observing the subsequent dynamics.

***Twin surrogate algorithm.*** First, we will explain the twin surrogate algorithm. This method shuffles an embedded vector sequence $\boldsymbol{v}(1), \ldots, \boldsymbol{v}(M)$ according to certain rules, where $M$ is the number of vectors. Here, $\boldsymbol{v}(i)$ is $\{z_i, z_{i-1}, \ldots, z_{i-(E-1)}\}$, and $E$ is an embedding dimension obtained from a simplex projection (6). Specifically, this method consisted of the following four steps:

1. For the $M$ vectors $\boldsymbol{v}(1), \ldots, \boldsymbol{v}(M)$ created by embedding the original time series data, create a recurrence matrix with $i, j = 1, \ldots, M$

$$R_{i,j} = \Theta(\delta - \|\boldsymbol{v}(i) - \boldsymbol{v}(j)\|)$$

   where $\Theta(\cdot)$ is a Heaviside function (i.e., $\Theta(\cdot) = 0$ for $\cdot < 0$ and $\Theta(\cdot) = 1$ for $\cdot \geq 0$) and $\|\boldsymbol{v}(i) - \boldsymbol{v}(j)\|$ denotes the maximum norm (the maximum absolute value of each component of $\boldsymbol{v}(i) - \boldsymbol{v}(j)$). This recurrence matrix indicates whether any two vectors $\boldsymbol{v}(i)$ and $\boldsymbol{v}(j)$ are farther from the threshold $\delta$ by 0 or 1. Therefore, the recurrence matrix contains the same information as the original nonlinear dynamical system attractor (7), although with less resolution.

2. If $R_{i,k} = R_{j,k}$ for all $k = 1, \ldots, M$, the pair $\boldsymbol{v}(i)$ and $\boldsymbol{v}(j)$ is called a twin. On a recurrence matrix of 0 and 1, it is impossible to distinguish between two vectors that are twins.

3. The $i$-th surrogate datum is denoted by $\boldsymbol{v_s}(i)$, and the original vector $\boldsymbol{v}$ is randomly selected as the first one, $\boldsymbol{v_s}(1) = \boldsymbol{v}$.

4. If $\boldsymbol{v_s}(j) = \boldsymbol{v}(m)$, then $\boldsymbol{v_s}(j+1) = \boldsymbol{v}(m+1)$ if $\boldsymbol{v}(m)$ does not have a twin. In contrast, if $\boldsymbol{v}(m)$ has $\boldsymbol{v}(n)$ as a twin, then $\boldsymbol{v_s}(j+1) = \boldsymbol{v}(m+1)$ or $\boldsymbol{v_s}(j+1) = \boldsymbol{v}(n+1)$ with equal probability, even if there are more than two twins.

Repeat Step 4 to create surrogate data with the same length as the original. The surrogate data shall be rejected if they reach the end of the original vector (i.e., $\boldsymbol{v}(M)$) before it reaches the same length as the original, and the end vector $\boldsymbol{v}(M)$ does not have a twin.

Although this method requires the parameter $\delta$, the dependence on $\delta$ has been shown to be small (5), and the best results seem to be obtained when the ratio of 1 in the recurrence matrix $R_{i,j}$ is $0.05 \sim 0.2$. For the analysis in this paper, we adopted a $\delta$ that produced a ratio of 1 in $R_{i,j} = 0.125$. It is known that the surrogate data created by this method preserve the properties of the original time series. In particular, mutual information, which is a nonlinear property, is known to be conserved by twin surrogates, but it is not conserved by other surrogate methods (5).

***Responses to perturbations.*** Furthermore, to examine responses to perturbations, we improved the procedure of determining the initial value in Step 3 in the following four cases:

- (a) The case of a small perturbation (corresponding to small $S$). We considered a new, original time series starting from a randomly chosen vector $\boldsymbol{v}(i)(1 \leq i \leq M - \tau)$, where $\tau$ is the time after the perturbation. We defined the initial vector of the perturbed time series to be $\boldsymbol{v_s}(1) = \boldsymbol{v}(i)$. Then, we repeated Step 4. Finally, the new, original time series was $\boldsymbol{v}(i), \ldots, \boldsymbol{v}(i+\tau)$, and the perturbed series was $\boldsymbol{v_s}(1), \ldots, \boldsymbol{v_s}(1+\tau)$. These two time series started from the same values, but they exhibited different dynamics via twins. Therefore, $\boldsymbol{v_s}(1), \ldots, \boldsymbol{v_s}(1+\tau)$ corresponded to a time series starting from a slightly different initial state.

- (b) The case of a large perturbation (corresponding to large $S$). The new, original time series was the same as in case (a), namely $\boldsymbol{v}(i), \ldots, \boldsymbol{v}(i+\tau)$. The initial value $\boldsymbol{v_s}(1)$ of the perturbed time series was randomly chosen from vectors that were in the top 5% of the distribution of $\|\boldsymbol{v}(i)\|$ where $\|\cdot\|$ means the Euclidean norm. Then, we repeated Step 4. The perturbed time series $\boldsymbol{v_s}(1), \ldots, \boldsymbol{v_s}(1+\tau)$ corresponded to a time series starting from a drastically changed initial state.

- (c) The case of stimuli that evoke straight movements (corresponding to small $S'$). To examine behavioral switching according to the type of stimulus, we randomly selected an initial value from vectors that were in the bottom 5% of the distribution of $\|\boldsymbol{v}(i)\|$. Then we generated subsequent time series with the twin surrogate method. Finally, we compared the positions constructed from $\boldsymbol{v_s}(1)$ and $\boldsymbol{v_s}(1+\tau)$.

- (d) The case of stimuli evoking turns (corresponding to large $S'$). We randomly selected an initial value from vectors that were in the top 5% of the distribution of $\|\boldsymbol{v}(i)\|$. Otherwise, this case was the same as case (c).

Note that if we could not create the perturbed time series due to the absence of twins in entire time series or a shortage of original time series, we did not use the time series for analysis. The final sample size was $n = 263$.

### References


1. F Bartumeus, MGE da Luz, GM Viswanathan, J Catalan, Animal search strategies: a quantitative random-walk analysis. *Ecology* **86**, 3078–3087 (2005).
2. A Pikovsky, J Kurths, M Rosenblum, J Kurths, *Synchronization: a universal concept in nonlinear sciences.* (Cambridge university press) Vol. 12, (2003).
3. RM May, Simple mathematical models with very complicated dynamics. *Nature* **261**, 459 (1976).

4. K Kaneko, Clustering, coding, switching, hierarchical ordering, and control in a network of chaotic elements. *Phys. D: Nonlinear Phenom.* **41**, 137–172 (1990).
5. M Thiel, MC Romano, J Kurths, M Rolfs, R Kliegl, Twin surrogates to test for complex synchronisation. *EPL (Europhysics Lett.* **75**, 535 (2006).
6. G Sugihara, RM May, Nonlinear forecasting as a way of distinguishing chaos from measurement error in time series. *Nature* **344**, 734–741 (1990).
7. M Thiel, MC Romano, J Kurths, How much information is contained in a recurrence plot? *Phys. Lett. A* **330**, 343–349 (2004).
